# Accessory proteins increase the efficiency of RNA editing by Arabidopsis chloroplast editosomes

**DOI:** 10.1101/2024.04.05.588343

**Authors:** Jose Lombana, Maureen R. Hanson, Stéphane Bentolila

## Abstract

RNA editing modifies cytidines to uridines in plant organelle transcripts so that their sequences differ from the ones predicted from the genomic DNA. This process, conserved across most land plants, involves a family of RNA-binding proteins that has significantly expanded, the pentatricopeptide repeat (PPR)-containing proteins. In angiosperms, PPR proteins are found in editosomes associated with accessory proteins. The exact function of these accessory proteins has been unclear. Bacterial co-expression of an angiosperm synthetic factor and different accessory proteins, RIP2, RIP9, ORRM1 demonstrates their essential role in editing of an RNA target. The presence of ORRM1 and RIP2 or ORRM1 and RIP9 in bacteria with the PPR factor results in a target editing extent of 80%, which is similar to what is observed *in planta*. Accessory proteins increase the affinity of the PPR factor for the target RNA, likely the explanation of their role in improving editing efficiency. RNA-seq analysis of bacterial transcriptome in samples expressing various combinations of accessory proteins with the synthetic factor identified a total of 34 off-target editing events. Investigation of their upstream sequences that are recognized and bound by the synthetic factor allows the optimization of future designs to improve the specificity of this programmable RNA-editing factor.

## INTRODUCTION

Among the post-transcriptional processes that influence gene expression, C-to-U RNA editing in plants exhibits several unique features. Editing in plants is restricted to genome-containing organelles—chloroplasts and mitochondria. Unlike other types of RNA editing, the purpose of plant RNA editing is not to generate multiple proteins from the same transcript; instead, its role is to rectify T-to-C mutations in critical locations of transcripts, ensuring the production of functional proteins (1). A typical vascular plant requires over 600 RNA editing-mediated corrections in its organelle transcripts (2). While previous research has shed some light on the editing mechanism, there is still much to be discovered—information that may eventually enable the biotechnological exploitation of this phenomenon.

How do plants achieve so many specific changes from Cs to Us? Through evolution, a modular family of RNA-binding proteins has significantly expanded, often comprising over 400 members in a single plant species (3). Pentatricopeptide Repeats (PPR), 35 amino-acid repeats presented in tandem, can bind RNA in a sequence-specific manner. Immediately upstream of C targets of plant organelle editing are short *cis*-elements to which the PPR repeats bind (4). Then a deaminase activity, either present at the C-terminus of the PPR protein (a ’DYW domain’) or recruited as a trans-factor, carries out the C-to-U conversion (5–7).

In vascular plants, the PPR proteins studied do not act alone, even if they carry their own deaminase activity. Instead, a complex set of trans-factors interacts with each PPR protein to form a small RNA/protein complex termed the editosome—typically 400 to 600 kD in size (8). A major question is WHY these accessory factors exist. There are a few known PPR-DYW proteins in a non-vascular plant—a moss—that can perform C-to-U editing without any accessory factors (9). One hypothesis is that plant accessory factors improve efficiency (percentage of the transcript population that is edited) and/or selectivity (editing only the important C target and not neighboring Cs or distant ones with similar *cis*-elements).

The identification of a synthetic angiosperm chloroplast editing factor active in planta and in *E. coli* (10), allows use of a bacterial system to investigate the role of additional accessory factors. The synthetic PPR, dsn3PLS-DYW, was designed with 3 PLS motifs to recognize *rpoA* C-200, which is not edited in Arabidopsis homozygous for the *clb19* mutation (11). A C-terminal DYW domain provides the deaminase catalytic activity. Dsn3PLS-DYW was able to restore a level of editing extent around 45% in the *clb19* mutant plant, but expression in *E. coli* resulted in only about 10% editing of an *rpoA* target sequence. However, co-expression of either the co-factor RIP2/MORF2 or RIP2/MORF9 significantly increased the level of editing to around 35% (10).

RIP2/MORF2 is only one of several accessory factors that are known to affect the efficiency of editing by CLB19 in Arabidopsis, according to mutant analysis (Table 1) (2,11–15). In this report, we investigated the role of additional accessory factors, either alone or in combination with each other, in the efficiency of editing of *rpoA*-C200 in *E. coli*. We tested whether accessory factors affected the affinity of the PPR protein for its target and demonstrated a correlation of improved affinity with increased editing extent. RNA-seq analysis of bacterial transcriptomes demonstrated that an increase in the efficiency of editing of the target by the accessory proteins was also correlated with an increase in off-target editing.

**Table 1.**
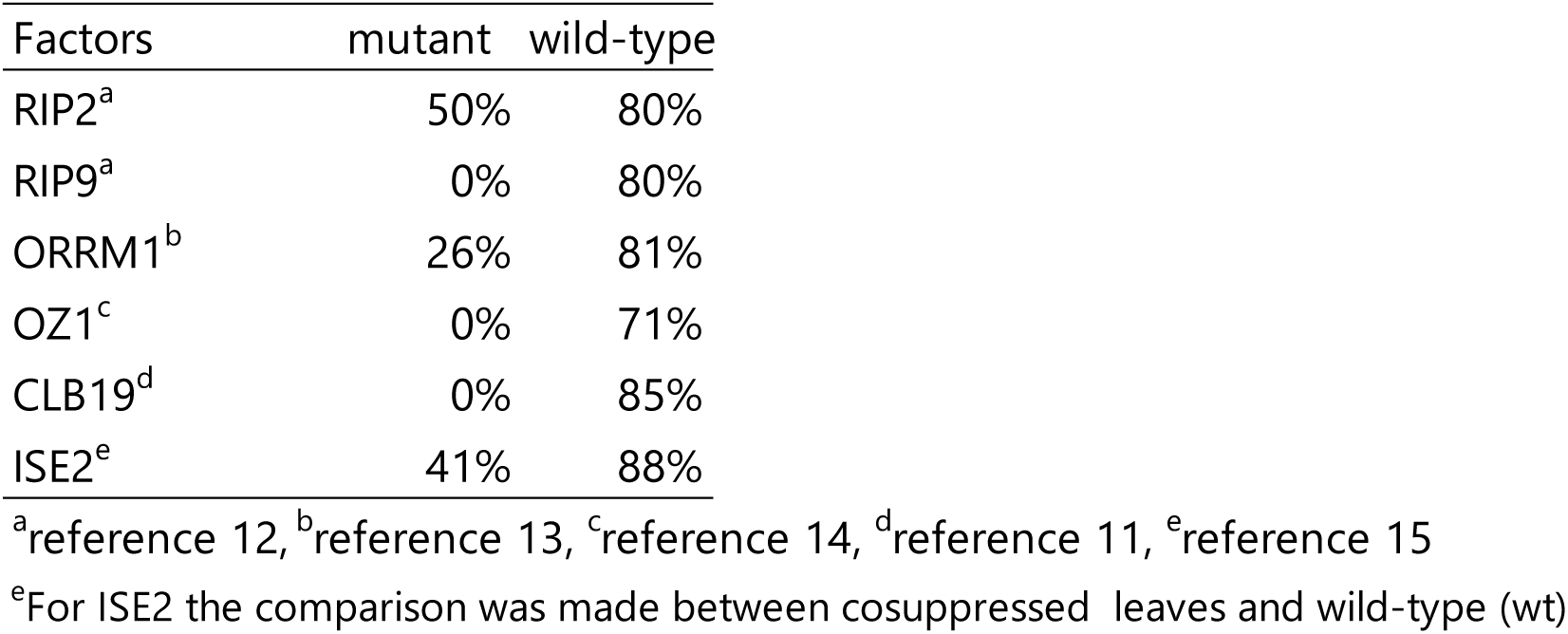
Editing extent of *rpoA* -C200 in mutant or silenced genes encoding accessory proteins.

| Factors | mutant | wild-type |
| --- | --- | --- |
| RIP2<sup>a</sup> | 50% | 80% |
| RIP9<sup>a</sup> | 0% | 80% |
| ORRM1<sup>b</sup> | 26% | 81% |
| OZ1<sup>c</sup> | 0% | 71% |
| CLB19<sup>d</sup> | 0% | 85% |
| ISE2<sup>e</sup> | 41% | 88% |

<sup>a</sup>reference 12, <sup>b</sup>reference 13, <sup>c</sup>reference 14, <sup>d</sup>reference 11, <sup>e</sup>reference 15
<sup>e</sup>For ISE2 the comparison was made between cosuppressed leaves and wild-type (wt)

## MATERIALS AND METHODS

### Bacterial strains

*Escherichia coli* strains used in this study were NEB 10-β competent cells (New England Biolabs, Ipswich, MA, USA, https://www.neb.com/en-us) for cloning and Rosetta 2 (DE3) (Novagen) for protein expression and *E. coli* RNA editing experiments.

### Plasmids

Duet vectors (Novagen) pETDuet-1, pCDFDuet-1 and pCOLADuet-1 are T7 promoter bacterial expression vectors designed to co-express two proteins in *E. coli*. These Duet vectors carry compatible replicons and antibiotic resistance markers and may be used together in appropriate host strains to co-express up to six proteins.

### Primers

The primers and oligonucleotides are listed in Supplementary Table S1 and were obtained from Integrated DNA Technologies (IDT, Coralville, IA, USA, https://www.idtdna.com)

### Cloning, bacterial expression

The synthetic factor nucleotide sequence was derived from the amino acid sequence published by Royan et al. (2021) and the corresponding DNA was synthesized by GenScript (GenScript, New Jersey, USA; https://www.genscript.com/) and cloned into the *NcoI* and *BamHI* sites of the pETDuet-1 vector. The accessory proteins encoding genes were obtained by PCR reaction with the primers listed in Supplementary Table S1, either directly from Arabidopsis genomic DNA (RIP2 and RIP9) or after RT-PCR from cDNA (ORRM1, OZ1, ISE2). The transit peptide-encoding sequences were predicted by using TargetP2 (https://services.healthtech.dtu.dk/services/TargetP-2.0/) and was removed from all the accessory protein encoding genes. After the PCR reaction the amplicons were cloned into the pCR™8/GW/TOPO vector (Thermo Fisher Scientific, Waltham, MA, USA; https://www.thermofisher.com) and the sequences verified to be accurate. Once the sequences were validated, the plasmids were digested with the appropriate restriction enzymes (See Supplementary Table S1 for details) run on agarose gel and the bands corresponding to the accessory proteins encoding DNA gel-purified and cloned into the expression vectors. The sequence of every expression vector used in this study was checked by whole plasmid sequencing performed by eurofinsgenomics (Louisville, KY, USA; https://eurofinsgenomics.com). Bacterial culture followed the protocol of Oldenkott et al. (2019); 1 ml samples were harvested, centrifuged at maximum speed for 10 minutes at 4°C, the pellets frozen in liquid nitrogen and stored at -80 °C until further use for protein-RNA analysis.

### Protein Extraction

Cell pellets were resuspended in 500 µL of Lysis Buffer (50 mM Tris-HCl, pH 8.0; 1 mM EDTA; 2 mM DTT). Cells were lysed using a 25-second sonication pulse (Heat Systems Ultrasonic Processor W-380, 70% output power, level 5 output control), ensuring complete disruption.

Protein concentrations were determined using the Bradford assay (Bio-Rad Catalog #500-0006), with concentrations normalized by diluting samples with Lysis Buffer to match the sample with the lowest protein concentration. Samples were then mixed with an equal volume of 2X Laemmli Buffer (Bio-Rad Catalog #161-0737), gently mixed to avoid frothing, heated in a 95°C water bath for 10 minutes, and then cooled on ice for 2 minutes. Samples were stored at -20°C for short-term and at -80°C for long-term storage.

### SDS-PAGE and Immunoblotting

10 µL of the normalized protein samples were loaded onto a Mini-PROTEAN TGX Any kD (Bio-Rad Catalog #456-9036) SDS-polyacrylamide gel and electrophoresed in 1X SDS-PAGE Running Buffer (25 mM Tris, 192 mM glycine, 0.1% w/v SDS) at 200 V for approximately 35 minutes. After electrophoresis, the gel was rinsed thoroughly with deionized water to remove any residual SDS. The gel was equilibrated in cold 1X Transfer Buffer (25 mM Tris, 192 mM glycine, 20% methanol) for 5 minutes at 4°C prior to transfer. Protein transfer was performed using a nitrocellulose membrane (0.45 µm, Thermo Scientific, REF.88018) in 1X Transfer Buffer using the Trans-Blot Cell (Bio-Rad, Serial No. 49BR32383) at 100 V for 1 hour at 4°C, ensuring the current did not exceed 1.5 A as recommended by the manufacturer. The membrane was dried on clean filter paper at 4°C for 20 minutes to affix the proteins. All subsequent steps were carried out in a black Western blot box (MTC Bio, Cat. No. B1200-7BK) to prevent fluorophore quenching. Blocking was performed at room temperature for 30 minutes using EveryBlot Blocking Buffer (Bio-Rad, Cat. #12010020). Primary antibody incubation was carried out overnight at 4°C in a 1:5,000 dilution of the primary antibodies in EveryBlot Blocking Buffer with gentle rocking. The membrane was washed twice with 1X TBST and twice with 1X TBS, each for 5 minutes with moderate rocking. Secondary antibody incubation was conducted at room temperature for 1 hour in a 1:10,000 dilution in EveryBlot Blocking Buffer, followed by two washes in 1X TBST and two in 1X TBS, each for 5 minutes with moderate rocking. The membrane was dried on clean filter paper at room temperature in the dark for at least 20 minutes before imaging. Imaging was performed using the LI-COR Odyssey Imaging System, employing DyLight 700, DyLight 800, and RGB channels for colorimetric ladder visualization. Primary Antibody Mix: (1:5,000): Rabbit anti-Stag pAb (Sino Biological #101290-T38) and Mouse anti-His-tag mAb (GeneScript Cat. No. A00186). Secondary Antibody Mix: (1:10,000): AlexaFluor 488 goat anti-rabbit IgG and AlexaFluor 546 goat anti-mouse IgG or Goat Anti-Mouse IgG (H + L) DyLight 680 Conjugated (Invitrogen REF 35518), Goat anti-Rabbit IgG (H&L) DyLight 800 Secondary Antibody (Invitrogen REF SA535571).

### Ni²⁺ Affinity Purification

A 1-L bacterial culture was initiated from a 10 mL fresh pre-culture. Upon reaching an optical density (OD₆₀₀) of 0.4 to 0.7, the culture was cooled on ice for 30 minutes and induced with IPTG to a final concentration of 50 µM. Concurrently, ZnSO₄ was added to a final concentration of 0.4 mM. The culture was then incubated at 16°C for 20 hours. Post incubation, the culture was harvested by centrifugation at 6,000 × g for 30 minutes at 4°C, and the supernatant was discarded. The cell pellet was flash-frozen in liquid nitrogen. The cell pellet was resuspended in a volume ten times its mass of lysis buffer (25 mM Tris-HCl, pH 7.2; 250 mM NaCl; 10% glycerol; 10 mM imidazole; 0.1% NP-40). Lysis was facilitated by the addition of lysozyme to a final concentration of 0.5 mg/mL, followed by 25 sonication pulses of 1 minute each, with 5 minutes of incubation on ice between pulses. The lysate was clarified by centrifugation at 16,639 × g for 30 minutes at 4°C, and the supernatant was then filtered through a 0.22 µm cellulose acetate filter using a vacuum system. The clarified lysate was applied to a 20-mL Econo-Pac polyethylene gravity column containing 1.5 mL of pre-equilibrated HisPur Ni²⁺ NTA agarose bead resin (Thermo Scientific, Prod# 88221) and allowed to bind for 30 minutes. The column was washed five times with 20 mL of wash buffer (25 mM Tris-HCl, pH 7.2, 22°C; 250 mM NaCl; 10% glycerol; 50 mM imidazole). Elution was performed by resuspending the resin in 3 to 5 mL of elution buffer (25 mM Tris-HCl, pH 7.2, 22°C; 250 mM NaCl; 10% glycerol; 200 to 400 mM imidazole) for 20 minutes. Eluted fractions were dialyzed against 500 mL of 20G buffer (25 mM Tris-HCl, pH 7.2, 22°C; 250 mM NaCl; 20% glycerol; 0.5 mM EDTA; 1 mM DTT) for 24 to 36 hours with gentle stirring at 4°C using a Side-A-Lyzer dialysis cassette (Thermo Scientific, Prod# 66380). The dialyzed protein was concentrated using Amicon 10K and 30K columns (Millipore, Ref. UFC803096; UFC801096) until the final protein concentration reached approximately 1 to 2 mg/mL. The protein was aliquoted, flash-frozen in liquid nitrogen, and stored at -80°C. The efficacy of the purification was assessed via Stain-Free SDS-PAGE using Mini-PROTEAN TGX Stain-Free Gels (BioRad, Cat.# 4568126) according to the manufacturer’s recommendations. The gels were imaged using the BioRad ChemiDoc MP Imaging System.

### Detection of RNA editing in *E.coli*

Total RNA was extracted from 1 mL of *E. coli* bacterial pellet using the Invitrogen™ PureLink™ RNA Mini Kit (Thermo Fisher Scientific, Waltham, MA, USA; https://www.thermofisher.com) according to the manufacturer’s instructions. Reverse transcription was performed on DNase-treated total RNA extract using Superscript™ III Reverse Transcriptase (Invitrogen) with the primers listed in Supplementary Table S1 according to the manufacturer’s instructions. RT-PCR Bulk sequencing of the *rpoA* -C200 target was performed to assay the editing extent. The editing extent was calculated from the electrophoretogram by using BEAT, a python program developed to quantify base editing from Sanger sequencing (16). For each accessory protein or combination of accessory proteins tested we performed three biological replicates.

### Statistical treatment of bacterial expression experiments

To examine the variability across the outcomes of 46 distinct experiments, each involving three independent measurements from biological replicates, a specific statistical framework was utilized. Analysis of Variance (ANOVA), conducted using the stats package (version 3.6.2) in R, was implemented to evaluate significant differences among the mean values of the experimental groups, revealing significant disparities (Pr < 2e-16, F = 303.1). Given the significant findings from the ANOVA, Tukey’s Honestly Significant Difference (TukeyHSD) test, also performed with the stats package (version 3.6.2) in R and was subsequently applied for pairwise comparisons among the experimental conditions.

### RNA electromobility shift assays (REMSA)

REMSAs were performed following the protocol of Royan et al. (2021). The fluorescence was visualized on an Amersham Typhoon system (GE Healthcare), with a filter for the Cy5 fluorophore and quantified with the analysis toolbox from the ImageQuantTL software version 8.2 (GE Healthcare).

### AlphaFold predictions

For the protein structure predictions of multiple sequences, including those from dsn3PLS-DYW (untagged) and the truncated variants of RIP2(89-186) and RIP9(86-192), ColabFold: AlphaFold2 integrated with MMseqs2 was employed (3). This advanced tool facilitated the generation of highly reliable structural predictions, as indicated by the precision metrics: predicted Local Distance Difference Test (pLDDT) and predicted Template Modeling (pTM) scores, alongside the interface predicted TM score (ipTM) for the complexes of dsn3PLS-DYW with RIP2(89-186) or RIP9(86-192). The accuracy of these predictions is further substantiated by Predicted of Aligned Error (PAE) graphs, provided in the supplementary materials (Supplementary Figure S7)

dsn3PLS-DYW + **RIP2**(**89-186**): pLDDT=93.1, pTM=0.885, ipTM=0.932
dsn3PLS-DYW + **RIP9**(**86-192**): pLDDT=93.1, pTM=0.878, ipTM=0.958

To analyze the structural alignment and visualization, the predicted models were superimposed using UCSF Chimera (developed by the Resource for Biocomputing, Visualization, and Informatics at the University of California, San Francisco, with support from NIH P41-GM103311, (17)), focusing on the chains corresponding to dsn3PLS-DYW. This approach facilitated a detailed comparison between the chains, which were pseudo-colored to enhance differentiation. Visualization of the structures, including the generation of images and movies, was also accomplished using UCSF Chimera, ensuring a comprehensive representation of the structural dynamics and alignments observed.

### Library construction for RNA-seq analysis

Total RNA was extracted from 1 mL of *E. coli* bacterial pellet as for the detection of RNA editing. RNA concentration and integrity were assessed using a Qubit 2.0 Fluorometer (Invitrogen) with the Qubit RNA BR Assay Kit (Invitrogen Ref No. Q10211) and an Agilent 2100 Bioanalyzer (Agilent Technologies) with the RNA 6000 Nano Kit (Agilent, Cat No. 5067-1511). RNA libraries were treated with the FastSelect rRNA depletion kit from Qiagen for bacterial depletion and a directional library preparation was used at Cornell RNA Core facility. The libraries were sequenced on an Illumina NovaSeq X instrument with unpaired-end 150 bp reads.

### Quality Control and Read Processing

Raw sequencing reads were assessed using FastQC (v0.12.1) to ensure data quality. Adapter sequences and low-quality bases were removed using Cutadapt (v4.9) in a two-step process: adapters were first removed based on the provided sequences, and then reads were hard-trimmed by 10 nucleotides from both 5’ and 3’ ends. Quality trimming was applied with a threshold of Q20.

A comprehensive reference genome was constructed by combining the E. coli reference genome (NCBI accession number: CP10816.1) with the pETDuet-rpoAC200 plasmid, containing the dsn3PLS-DYW synthetic PPR protein gene and the rpoAC200 target sequence. The pRARE2 plasmid (included with Rosetta cells and containing rare codon genes and the chloramphenicol resistance cassette) was sequenced and assembled using long-read sequencing services provided by Eurofins Genomics. This plasmid was included in the reference genome for alignment purposes but was excluded from the final off-target analysis due to the absence of appropriate annotations.

### Alignment and BAM File Processing

Sequencing reads were aligned to the constructed reference genome using the BWA-MEM algorithm (bwa-mem2 v2.2.1). Default parameters were applied, including a minimum seed length (-k 19), band width for the banded alignment (-w 100), off-diagonal X-dropoff (-d 100), internal seed search parameter (-r 1.5), clipping penalty (-y 20 and -L 5), minimum chain length discard threshold (-c 500), and scoring metrics including match score (-A 1), mismatch penalty (-B 4), gap open (-O 6), and gap extension (-E 1) penalties, alongside the penalty for unpaired read pairs (-U 17). BWA-MEM’s intrinsic soft clipping was utilized to handle read alignments, particularly at read ends where mismatches often occur due to sequencing errors or low-quality.

BAM files generated by BWA-MEM were sorted and indexed using Samtools (v1.18). The samtools calmd function was employed to generate MD tags, preparing the alignments for variant calling with JACUSA2 (v2.0.4) (18). Technical duplicates were merged before alignment to ensure comprehensive coverage. Quality assurance of alignments was performed with Qualimap (v2.2.1), and HTML reports were generated for each sample.

### Identification of Off-Target RNA Editing Events

Off-target RNA editing events were identified using JACUSA2 (v2.0.4) with parameters optimized for detection accuracy (18). Call-2 flag was utilized to set comparisons between the RNA samples and the genomic DNA sequences. Libraries were analyzed in a strand-specific manner using the -P RF-FIRSTSTRAND option to maintain correct strand orientation. A base quality filter threshold (-q 20) was applied to exclude bases with a Phred score below 20 and reads with more than two mismatches (-filterNM 2) were excluded to enhance specificity. These settings were chosen to balance sensitivity and specificity, ensuring the identification of true RNA editing events while minimizing false positives.

VCF files generated by JACUSA2 were further analyzed in R (v4.1.1) using custom scripts. The vcfR package (v1.14.0) was used for parsing, followed by stringent filtering criteria. C->T and G->A transitions were selected based on the following: (1) a minimum editing fraction difference of 1% between DNA control and RNA samples, with an error rate limited to 1% of the percentage editing; (2) events supported by read depths above the 1% lower quantile; and (3) a JACUSA2 score threshold above the 95th percentile for samples with editing rates ≥10% and the 98th percentile for samples with editing rates <10%. These criteria ensured the exclusion of SNPs and sequencing errors, focusing on genuine C-to-U RNA editing events.

### Validation and Downstream Analysis

To eliminate sequencing or mapping errors and potential SNPs, we excluded common events between Rosetta samples without PPR or accessory proteins using a custom Python script FASTA sequences of potential off-targets were extracted using a custom Python program for further analysis with PPRmatcher (https://github.com/ian-small/PPRmatcher) and for generating RNA logos (WebLogo 2.8.2). The PPRmatcher (https://github.com/ian-small/PPRmatcher; dsn3PLS.motifs.txt; scoring tables = Kobayashi.tsv) JULIA script was modified to return scores for individual PPR-binding sites, providing granular insights into binding affinities.

Custom Python and R scripts were used to generate RNA heat maps based on the PPRmatcher scores, the RNA logos using WebLogo (https://github.com/WebLogo/weblogo), and the comparison tables, offering visual representations of RNA editing events and binding site distributions across all samples.

## RESULTS

### Simultaneous expression of RIP2, RIP9, or ORRM1 with dsn3PLS-DYW in *E. coli*

In order to express the PPR specificity factor along with individual accessory factors, we used the Duet vectors system (Novagen) that was developed to co-express multiple genes in *E. coli*. This system consists of five vectors, each of which is capable of co-expressing two proteins. Maintenance of four plasmids in a single bacterial cell is possible because of compatible replicons and drug resistance genes that these vectors harbor. pETDuet-1 carries the ColE1 replicon and bla gene (ampR) and was used to express the dsn3PLS-DYW, which was synthesized by GenScript (Figure 1). The *rpoA*-C200 target sequence that we used consisted of 39 nt, 33 nt upstream of the targeted C and 5nt downstream, and was cloned downstream of the dsn3PLS-DYW sequence. Upon transcription, both dsn3PLS-DYW and *rpoA*-C200 are carried on the same transcript. The accessory proteins were cloned in pCDFDuet-1, which carries the CloDF13 replicon and the *aadA* gene (streptomycin/spectinomycin resistance) and in the kanamycin-resistant pCOLADuet-1, which carries the ColA replicon. Each Duet vector has two T7lac promoters, two multiple cloning (MCS) regions, and a single T7 terminator for the cloning and expression of two open reading frames (ORFs). The first MCS encodes an amino-terminal 6-amino acid (aa) His-tag sequence for detection and purification, while the second MCS allows the fusion of an optional carboxy-terminal 15-aa S-tag sequence for detection, purification, and quantification. Each of the 5 accessory proteins known to affect editing extent in planta, RIP2, RIP9, ORRM1, OZ1, and ISE2 (Table 1) was cloned either in pCDF Duet-1 or pCOLA Duet-1 with a His-N tag, a C-S tag, or no tag (Figure 1).

**Figure 1:**
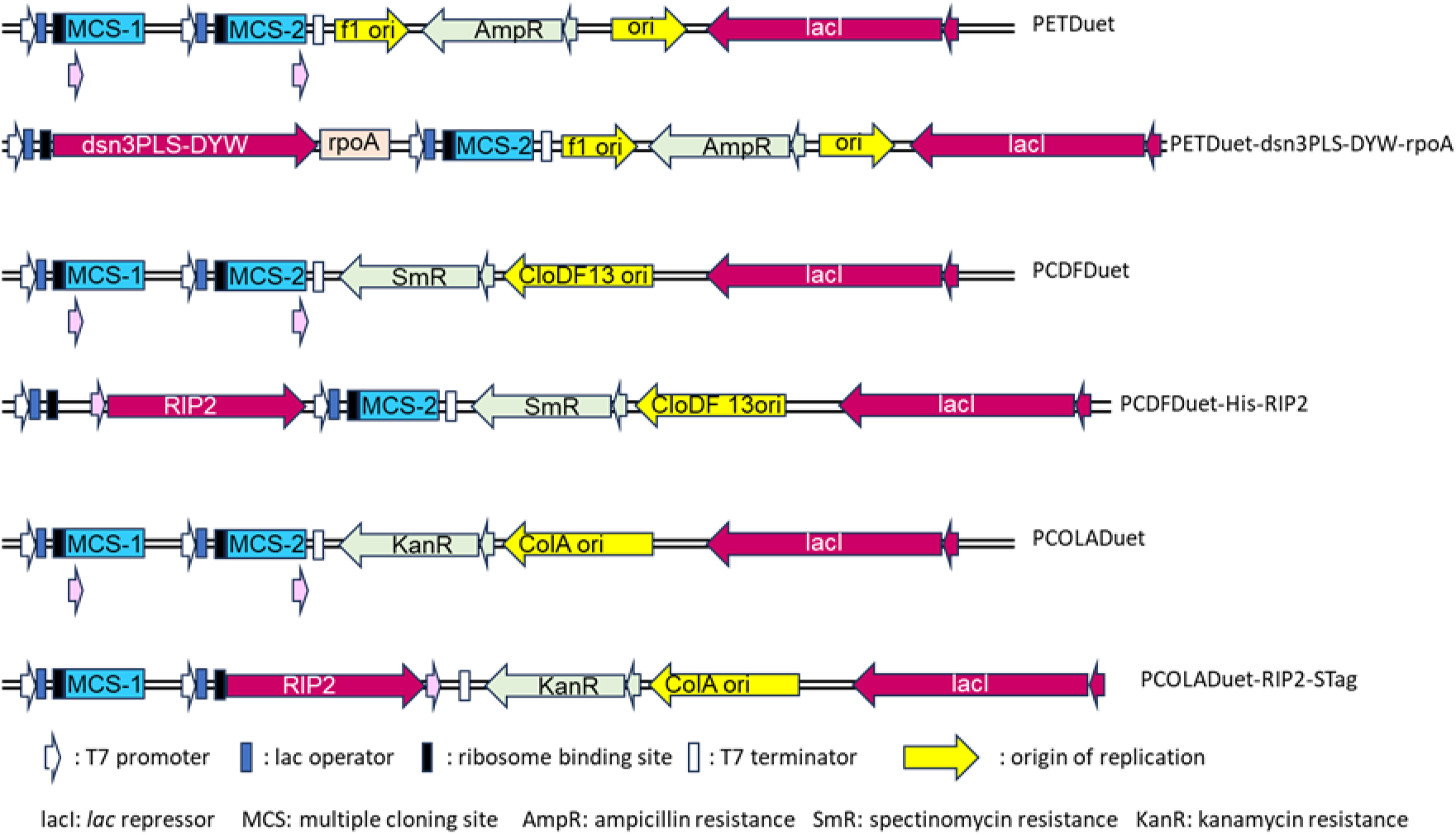
Simplified diagram of the vectors used. PETDuet, pCDFDuet and pCOLADuet are Duet vectors that are designed to allow the co-expression of multiple genes. The Duet vectors are T7 promoter expression vectors, each designed to co-express two proteins in *E. coli*. The Duet vectors carry compatible replicons and antibiotic resistance markers and may be used together in appropriate host strains to co-express up to six proteins. Below each vector is a representation of the vector with the gene it will express, pETDuet with the synthetic factor dsn3PLS-DYW, pCDFDuet or pCOLADuet with RIP2.

Although Royan et al. (2021) observed a low level of editing extent (5-10%) when the synthetic dsn3PLS-DYW factor was expressed by itself, we did not observe any editing extent in our system by expressing only the PPR editing factor (Figure 2). This difference could be due to our use of a different expression vector (pETM20 vs pETDuet-1) and thus a possible difference in the level of expression of the synthetic factor.

**Figure 2.**
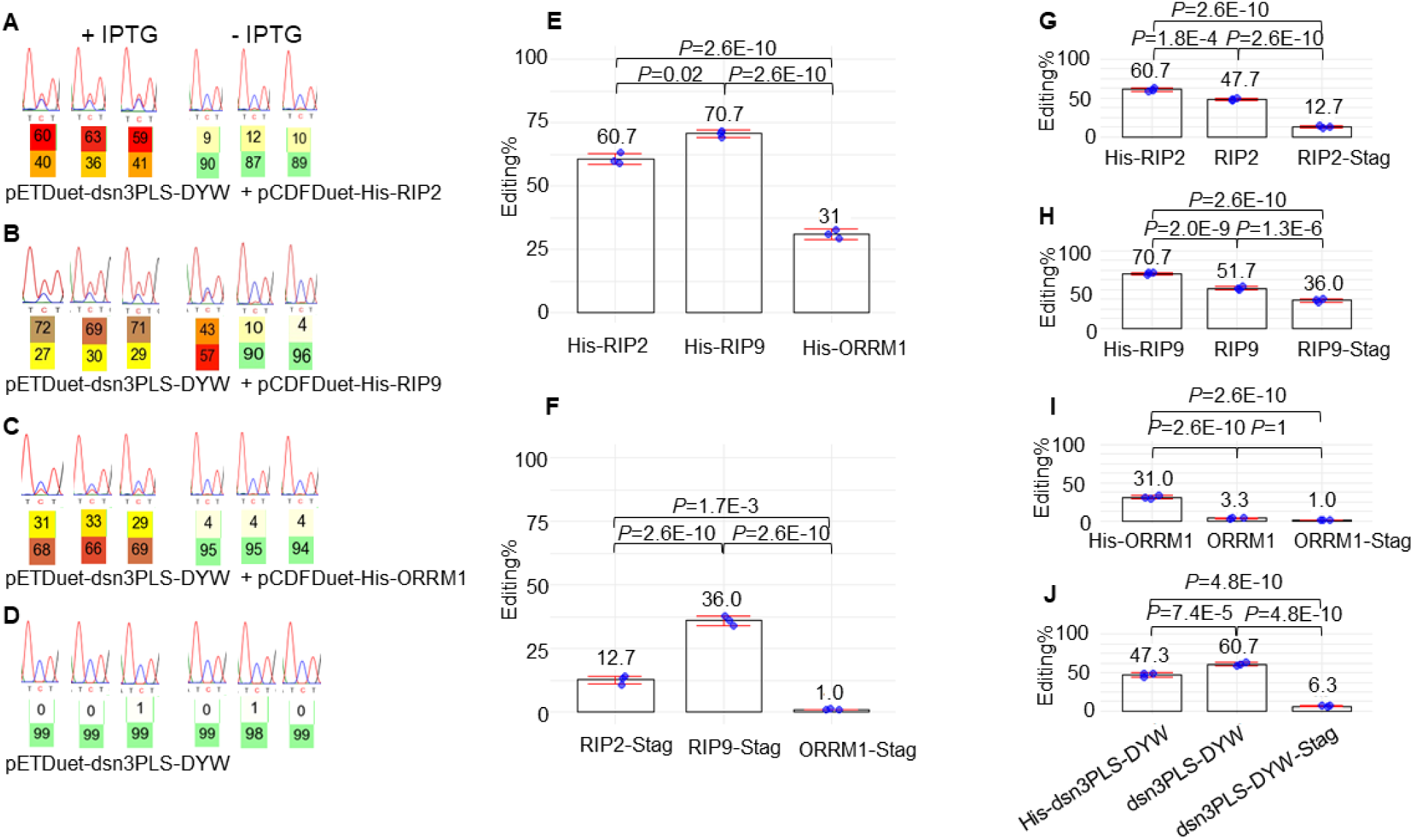
The accessory protein RIP9 is the most efficient in increasing the editing extent of *rpoA*-C200 by dns3PLS-DYW *in E. coli*. Left panel, electrophoretogram of RT-PCR bulk sequencing of the *rpoA*-C200 target when the synthetic factor is co-expressed with RIP2 (A), RIP9 (B), ORRM1 (C), or by itself (D). On the left (+IPTG) is the editing observed after induction by IPTG, on the right without induction. The C target is in the middle of the electrophoretogram showing 3 nucleotides T C/T T. Below the targeted C is the percentage of T (upper number) versus C (lower number) as computed by the BEAT software. The expression system is leaky as residual editing extent is observed even in the absence of induction. This residual editing in the absence of IPTG is more pronounced for efficient accessory protein RIP9 than for ORRM1. However, there is absolutely no editing when the synthetic factor is expressed by itself (D). Middle panel, with a tag either at the N terminus (His-Tag) (E) or at the C terminus (S Tag) (F) RIP9 is the most efficient accessory protein to increase the editing extent of the *rpoA*-C200 target. The editing extent of *rpoA*-C200 is given for three biological replicates in the presence of the synthetic factor and one accessory protein, RIP2 (left), RIP9 (middle) or ORRM1 (right). Each of the accessory protein was expressed using the pCDF-Duet1 vector. Right panel, RIP2 and ORRM1 are very sensitive to the presence of a tag at their C terminus while RIP9 is less affected. The addition of a tag at the N terminus is beneficiary to the editing function of the accessory proteins, particularly for ORRM1 (I). The editing extent of *rpoA*-C200 is given for three biological replicates in the presence of the synthetic factor and one accessory protein, RIP2 (G), RIP9 (H) or ORRM1 (I). Each of the accessory proteins was expressed using the pCDF-Duet1 vector. His tag is at the N terminus of the accessory protein, S tag at the C terminus. In J dsn3PLS-DYW was co-expressed with pCDF-His-RIP2.

When we expressed the synthetic PPR editing factor with RIP2, we observed approximately 60% editing, while RIP9 conferred 70% editing (Figure 2A, B). In contrast, Royan et al. (2021) observed only 35% editing. The level of editing extent of *rpoA*-C200 in our experiments was around 30% when the synthetic factor was co-expressed with ORRM1 (Figure 2C). These results were obtained with a His-tag attached at the N terminus of the accessory proteins (Figure 2E). We also tested these accessory proteins in a different configuration, with a S-tag attached at their C terminus (Figure 2F). Again, RIP9 was the most efficient accessory protein in increasing the editing extent of the *rpoA*-C200 target in the presence of the synthetic factor. However, the presence of a tag at the C terminus was detrimental to the function of the accessory proteins, as they all showed a significant reduction in their effect on the editing extent of the target. In the case of ORRM1, there was a complete obliteration of its function, with an editing extent that was close to zero (Figure 2F). Given their small sizes, 6 aa for the His-tag and 15 aa for the S-tag, the effect is likely caused by the position of the tag disrupting protein function.

Based on this observation, we tested the function of these three accessory proteins in the absence of any tag (Figures 2G, H, I). Surprisingly, a His-tag at the N terminus results in a higher editing extent for all the accessory proteins when compared to an absence of tag, possibly due to an increase in the stability of these proteins. This observation is particularly true for ORRM1 which, when deprived of a His-tag at the N terminus, experienced a severe reduction in the editing extent (3% vs 31%) of the target when co-expressed with dsn3PLS-DYW (Figure 2I).

Furthermore, we tested the position of the tag on the function of the synthetic factor itself. Unlike the accessory proteins, a His-tag at the N terminus of the dsn3PLS-DYW reduces the editing extent of the target when co-expressed with RIP2 (Figure 2J). Similarly to the accessory proteins, the addition of a S tag at the C terminus of the synthetic factor is very detrimental to its function, as the level of editing extent in the presence of RIP2 is severely reduced (Figure 2J).

Even though pCDF-Duet1 and pCOLA-Duet1 are supposed to have the same copy number (20-40) according to the Novagen user protocol, our experience in isolating these plasmids has generally been a low yield for pCOLA-Duet1. The lower copy number is reflected in a reduction of the expression of the accessory proteins in pCOLA-Duet1 when compared to pCDF-Duet1. The lower copy number and expression levels can affect the level of editing extent for certain accessory proteins in certain configurations, but not in others. For instance, a strain expressing ORRM1 with a His-tag experiences a reduction of almost half of the editing extent (17% vs 31%) depending on the expression vector used (pCOLA-Duet1 vs pCDF-Duet1), respectively (Supplementary Figure S1). A similar reduction occurs when RIP9 has either a His- or an S-tag, 71% vs 58%% of editing when expressed in pCDF-Duet1 vs pCOLA-Duet1, or 36%% vs 11%, respectively (Supplementary FigureS1). However, a strain expressing RIP2 with a His tag is not affected by the choice of expression vector; either results in a similar level of editing extent.

Although both OZ1 and ISE2 affect editing of *rpoA*-C200 in plant, neither of them affected editing when co-expressed by themselves with the synthetic factor (Supplementary Figure S2).

### ORRM1 and RIP2 (RIP9) act additively to contribute to the editing of *rpoA*-C200 by dsn-3PLS-DYW

In order to probe the role of ORRM1 in the presence of either RIP2 and RIP9, we tested the three combinations of two accessory proteins, RIP2 + RIP9, ORRM1 + RIP9, and ORRM1 + RIP2. Of these, ORRM1 +RIP2 or ORRM1 + RIP9 showed a very significant increase of the *rpoA*-C200 editing extent when compared to the individual contribution of each accessory protein (Figure 3). The editing extent in the presence of RIP2 and ORRM1 is almost the summation of each individual effect (82% vs 31% + 56%) suggesting that RIP2 and ORRM1 are acting in independent ways to allow the editing of *rpoA*-C200 by the synthetic factor (Figure 3A). A similar observation can be made for the combined effect of ORRM1 and RIP9 (Figures 3B, 3C). The editing extent in the presence of both ORRM1 and RIP9 is almost the summation of their individual effects. We tested RIP9 in both configuration with either a N-His tag (Figure 3B) or a C-S tag (Figure 3C): both experiments lead to the same conclusion on the cumulative effect of RIP9 and ORRM1 when they are co-expressed.

**Figure 3.**
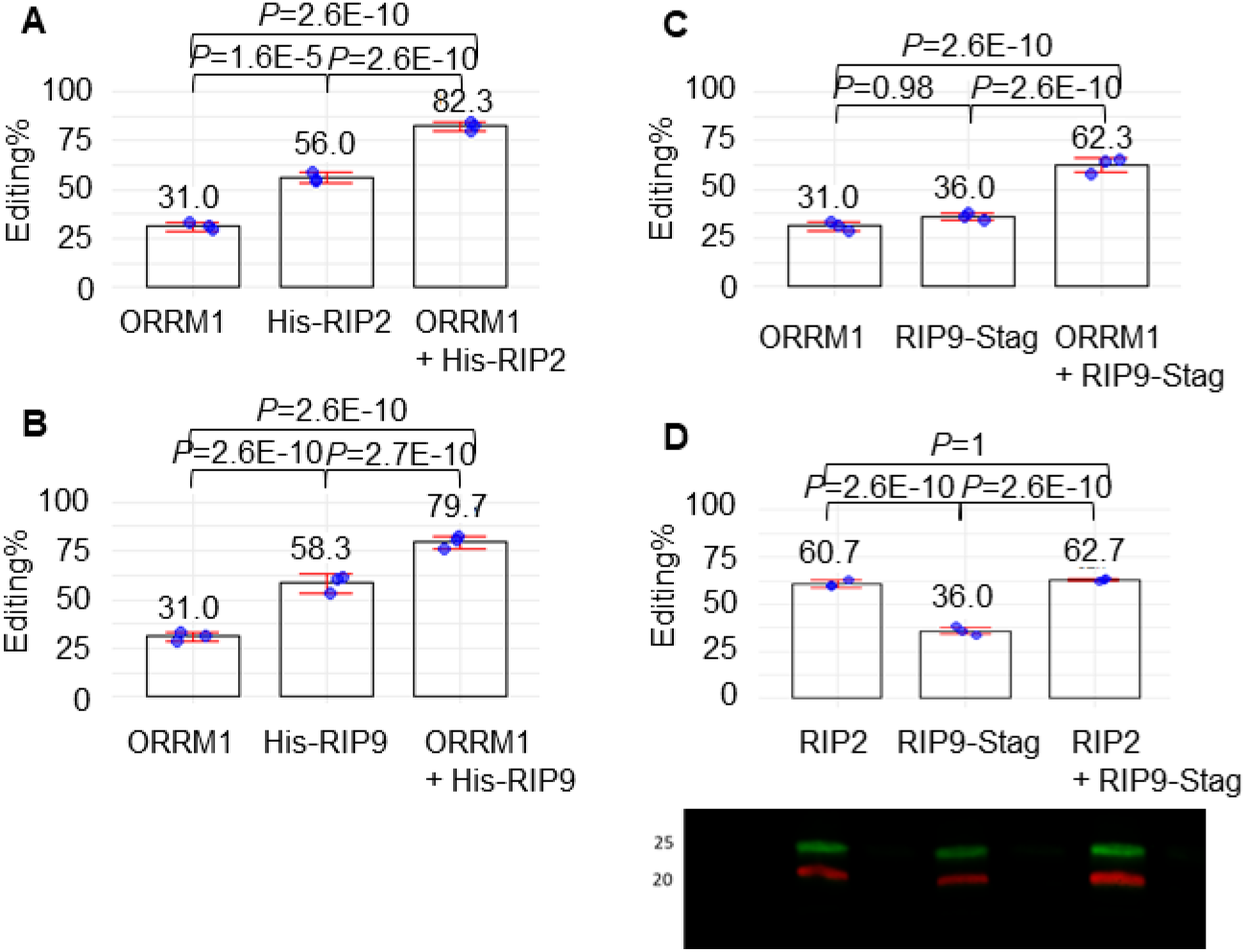
ORRM1 and RIP2 (RIP9) have an additive effect on the editing extent of the *rpoA*-C200 target. (A) Editing extent of the *rpoA*-C200 by dns3PLS-DYW in the presence of ORRM1 (left), RIP2 (middle) or both ORRM1 and RIP2 (right). (B) and (C) Editing extent of the *rpoA*-C200 by dns3PLS-DYW in the presence of ORRM1 (left), RIP9 (middle) or both ORRM1 and RIP9 (right). ORRM1 was expressed in pCDF-Duet1 with a His-tag, and RIP2 and RIP9 were expressed in pCOLA-Duet1 (A, B, C). (D) RIP2 and RIP9 were expressed in pCDF-Duet1. Below the graph is shown a Western blot demonstrating that both RIP2 (red) and RIP9 (green) are expressed in the three biological replicates (on the left are indicated the maker size in kDa). The editing extent of *rpoA*-C200 is given for three biological replicates.

In contrast, RIP2 and RIP9 do not show any sign of cooperation when they are co-expressed in the presence of the synthetic factor (Figure 3). The level of editing extent of the target is not significantly different in the presence of RIP2 and RIP9 (63%) from the editing extent when only RIP2 is present (61%).

### Co-expression of OZ1 or ISE2 with RIP2 does not increase editing efficiency

As described above, OZ1 and ISE2 do not affect the editing extent of the target when they are co-expressed with the synthetic factor (Supplementary Figure S2). In order to determine the roles of OZ1 and ISE2 in editing of *rpoA*-C200, we co-expressed each one in the presence of RIP2 and the synthetic PPR editing factor. The editing of the target is reduced when OZ1 with a His tag is co-expressed with RIP2 when compared to RIP2 alone (52 % vs 61%, Supplementary Figure S3A), indicating an inhibitory effect of His-OZ1. Therefore, we co-expressed RIP2 and OZ1 without a tag and found that the editing extent was slightly increased when compared to RIP2 alone (65% vs 56%, Supplementary Figure S3B). Both the inhibitory and stimulatory effects exhibit a *P* value of 0.08.

### RIP2 and RIP9 increase the affinity of the synthetic factor for its RNA ligand, while ORRM1 increases the affinity only when combined with another accessory protein

How do the accessory proteins increase the editing efficiency of the synthetic dsn3PLS-DYW factor for its target in bacteria? The RNA binding activity of a designer PLS-type PPR protein was reported to be drastically increased on RIP9 (also referred to as MORF9) binding via conformational changes of the PPR protein (19). We used gel-shift assays to determine whether RIP2, RIP9, or ORRM1 could increase the affinity of dsn3PLS-DYW for its target. Dsn3PLS-DYW was cloned in the expression vector PETM20 and was fused at its N-terminus with thioredoxin (109aa) followed by a 6xHis tag. As a result, the recombinant dsn3PLS-DYW carried a 133 aa tag at its N-terminus (Supplementary Figure S4). RIP2, RIP9 and ORRM1 were expressed from the pCDF-Duet1 vector with a His tag. We increased the length of the tag to 10xHis for RIP2 to improve its purification. Each recombinant protein was purified from overnight IPTG-induced cultures using Ni2+ affinity purification (Supplementary Figure S5).

The synthetic PPR editing factor can shift the RNA target, but with a limited efficiency (Figure 4A). We then tested ORRM1, RIP2, or RIP9 by adding an increasing amount of the accessory protein in the presence of a constant amount of the synthetic factor and the RNA target. No increase in the amount of the higher molecular mass complex was observed with ORRM1, demonstrating that ORRM1 has no effect on the RNA binding activity of the synthetic factor (Figure 4B). In contrast, there was a very noticeable increase in the complex when we repeated this experiment with RIP2 (Figure 4C) or RIP9 (Figure 4D). Both proteins are not able to bind the RNA target (no dsn3PLS-DYW lane in Figures 4C, D) by themselves, confirming that their effect is mediated by binding to the synthetic factor. The experiments presented in Figure 4 were repeated and the intensity of the lower band (unbound) and the upper band (bound) were quantified for each lane. Binding was estimated by plotting the fraction bound against the concentration of the accessory protein (Figure 4E). The estimated Kd for the synthetic factor was significantly reduced by the addition of RIP2 or RIP9. The enhancement of the RNA binding activity was more prominent with RIP9 than with RIP2 (Figure 4E).

**Figure 4.**
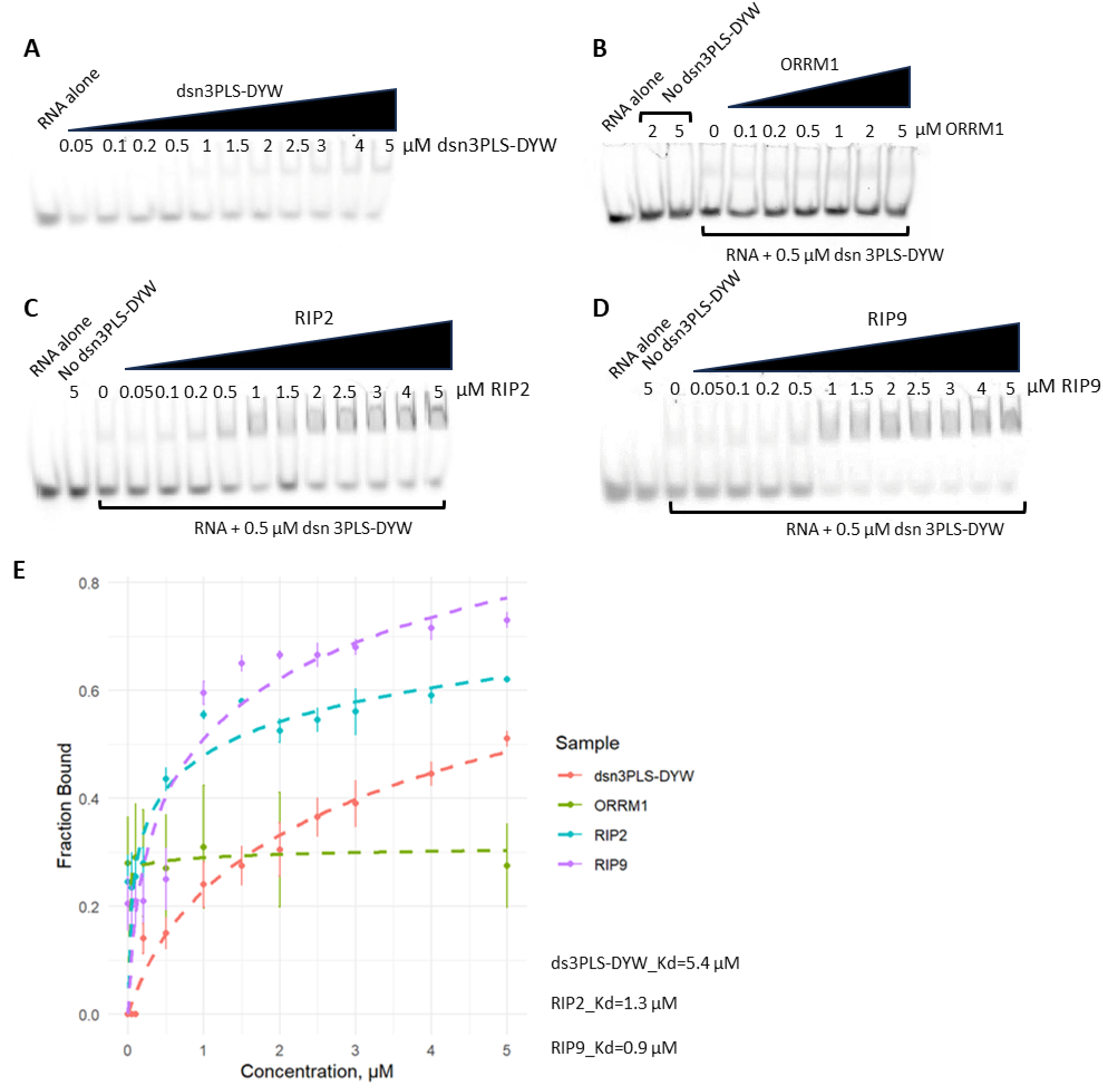
RIP2 and RIP9 but not ORRM1 increase the binding affinity of dsn3PLS-DYW protein for its RNA ligand. Gel shift assays were performed with dsn3PLS-DYW alone (A), or dsn3PLS-DYW in the presence of an increasing amount of ORRM1 (B), RIP2 (C) or RIP9 (D). RIP2 and RIP9 do not bind the RNA target by themselves (No dsn3PLS-DYW lane in C and D). (E) Fraction bound in each gel shift assay was plotted against the concentration of the accessory protein. Each reaction was duplicated. A logarithmic fitting model allowed to estimate the Kd for dsn3PLS-DYW and RIP2 and RIP9 with dsn3PLS-DYW.

Given that ORRM1, when co-expressed in bacteria with either RIP2 or RIP9, increases the editing efficiency of the target (Figure 3), we tested whether the enhanced binding activity of the synthetic factor by RIP2 could be influenced by the presence of ORRM1. Gel shift assays were performed with an increasing amount of RIP2, but with a constant amount of ORRM1. The amount of ORRM1 was serially increased from 0 µM to 4 µM (Figure 5A). The highest amount of ORRM1 tested (4 µM) resulted in a larger fraction bound than in the absence of ORRM1, particularly for the points corresponding to 0.2 µM, 0.5 µM, and 1 µM of RIP2 (compare purple and orange curve in Figure 5B). The affinity curve obtained with intermediate amounts of ORRM1 (0.5 µM and 1 µM) were between the ones obtained with 0 µM and 4 µM (Figure 5B).

**Figure 5.**
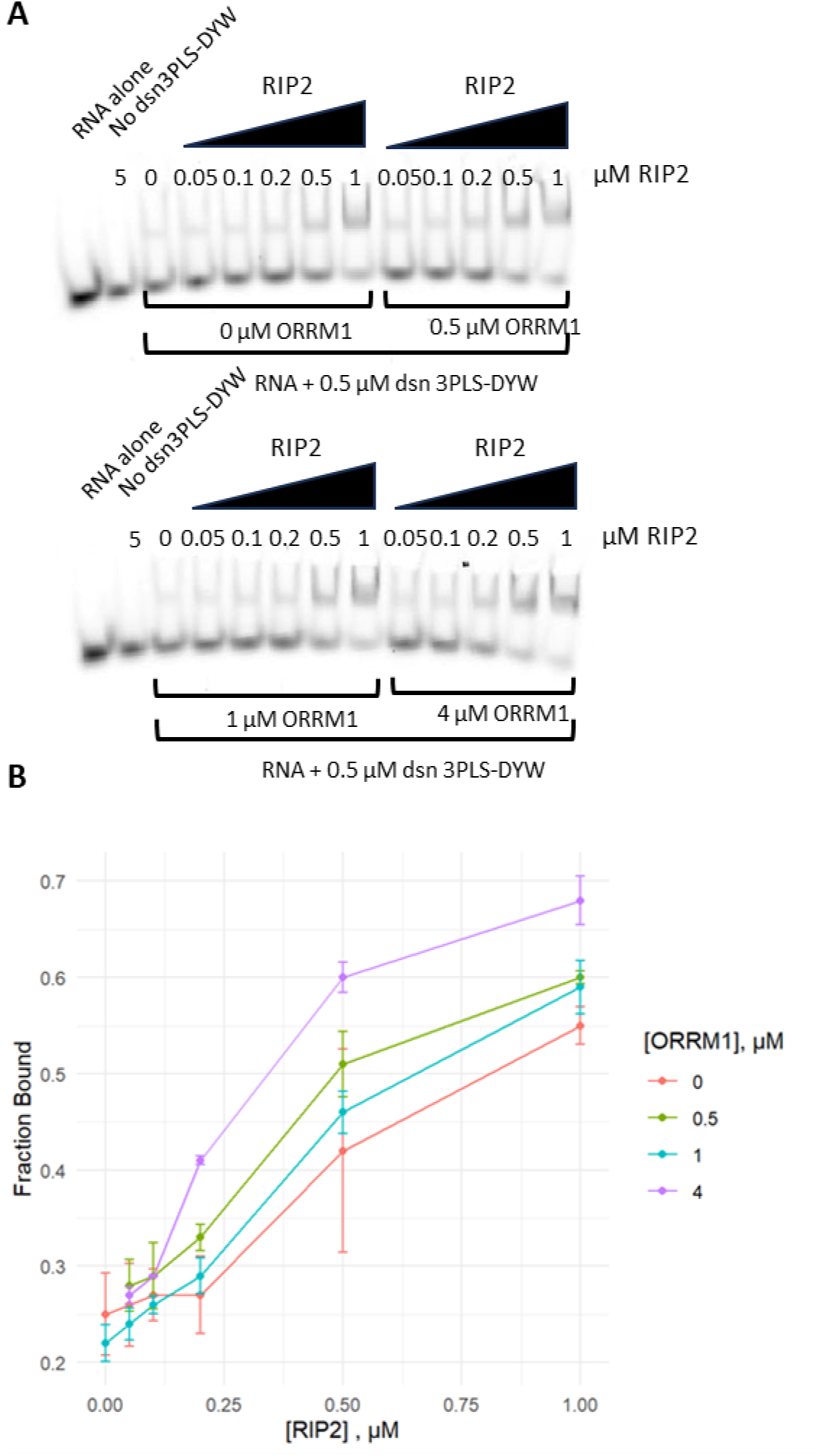
ORRM1 in the presence of RIP2 enhances the binding affinity of dsn3PLS-DYW protein for its RNA ligand. (A) Gel shift assays were performed with dsn3PLS-DYW in the presence of an increasing amount of RIP2 and a constant amount of ORRM1 : 0 µM (left upper gel), 0.5 µM (right upper gel), 1 µM (left lower gel), and 4 µM (right, lower gel). RIP2 does not bind the RNA target by itself (No dsn3PLS-DYW lane). (B) Fraction bound in each gel shift assay was plotted against the concentration of RIP2. Each reaction was duplicated. The fraction bound is more important for the higher amount of ORRM1 (4 µM, purple line).

Therefore, this experiment demonstrates that ORRM1 has a positive effect on the enhanced binding activity of the synthetic factor mediated by RIP2.

### RNA-seq analysis identifies 34 off-targets in the bacterial transcriptome

RNA-seq was performed on 14 bacterial RNA samples, 11 samples from bacteria expressing the synthetic factor with various combinations of accessory proteins, and 3 control samples: bacteria expressing the synthetic factor alone and bacteria not transformed, induced or not induced with IPTG (Supplementary Table S2). Editing of the target was identified in the 12 samples expressing the synthetic factor with its target. Editing of the target was 4.5% in the sample expressing only the synthetic factor, while no editing was detected by the less sensitive method of RT-PCR bulk sequencing (Figure 2D). Among the 12 samples expressing the synthetic factor, four did not show any off-target editing. In addition to the bacteria expressing only the synthetic factor, samples expressing ORRM1 (either with pCDF or pCOLA) and ORRM1 in the presence of ISE2 did not exhibit any off-target editing. The number of off-target editing events ranges from 4 to a maximum of 19, which was observed for sample 6, which co-expresses the synthetic factor with ORRM1 and RIP2 (Figure 6A).

**Figure 6.**
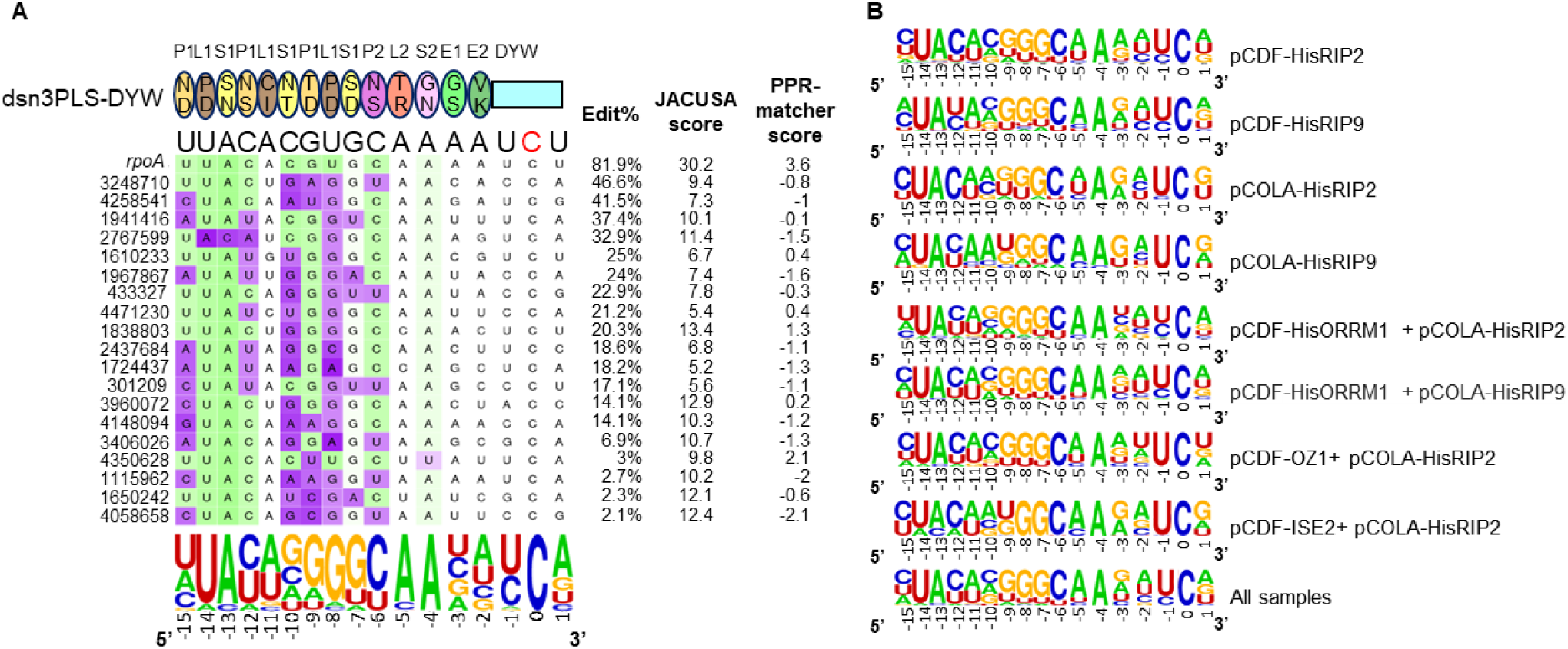
Identification of off targets reveals several positions in their sequences that deviate from the PPR recognition code. (A) Alignment of the synthetic editing factor with the sites edited in the bacterial transcriptome of Rosetta co-expressing dsn3PLS-DYW, the synthetic factor and the *rpoA* target (pETDuet) with ORRM1 (pCDF) and RIP2 (pCOLA). The dsn3PLS-DYW protein is represented by a model with oval for each PPR motif including the 5^th^ (upper aa) and last (lower aa) specificity-determining amino acids. Below the protein model is the sequence of the *rpoA* target site (the targeted C is in red). The coordinates of each off-target site on the bacterial chromosome are given on the left. The target sites are colored according to the PPR matcher score at each position, from favored (green) to neutral (white) to disfavored (purple). The percentage of edited transcripts at each site is indicated on the right. The JACUSA score reflects the confidence of a true editing event (the higher, the more confidence, see materials and methods for the threshold used). The PPR matcher score is a metric that indicates how well the sequence fits the PPR recognition model. The sequence logo indicates the nucleotide biases at each position and were computed with WebLogo excluding the *rpoA* target sequence and weighing each sequence with editing extent. (B) Sequence logos representing the consensus of off-targets for each sample. Like in (A) the *rpoA* target sequence was excluded and the sequences were weighed with editing extent. The logo at the bottom represents the consensus sequence of off targets found across all samples.

Several positions in the sequence of off-targets deviate from the expected predicted nucleotide based on the recognition code. These positions are highlighted in purple in Figure 6A. The most deviant position is found at -8 where a U is expected to be found but where predominantly a G is found in 15 of the 18 off-target sequences (Figure 6A). This bias occurs in all the samples analyzed as reflected by the logo sequence obtained for each of them and across all samples (Figure 6B). The next position to diverge significantly from the expectation based on the recognition code is -15, where a C is predominantly observed instead of a U in 6 of the 8 samples (Figure 6B). These two positions, -15 and -8, are the only ones that deviate consistently across all samples and show the same bias towards C and G, respectively (Figure 6B). Other positions exhibiting a different nucleotide in the consensus logo for some samples but not across all samples are -10, -9 and -12. At -10, 4 samples show a nucleotide different from the expected C while two samples and one sample show in their logo sequence a nucleotide different from the expected one at -9 and -12, respectively (Figure 6B).

It is noteworthy that the most deviant positions in the off-target sequences are observed where a pyrimidine is located, U at -15 and -8 and C at -10. The predictive value of the PPR matcher score for the level of editing extent of the off-targets is rather poor, as illustrated by the low level of editing, 3%, of the site located at 4350628, which has the highest score, 2.1, among the off-targets observed for sample 6, which co-expresses the synthetic factor with ORRM1 and RIP2 (Figure 6A). A significant, albeit weak, correlation (R^2^=0.12) between the editing extent of the off-targets and the PPR matcher score was only found for sample 1, which co-expresses the synthetic factor with pCDF-RIP2. Although the recognition code predicts a neutral or slightly favored nucleotide at position -5 and -4, respectively, we observed a very strong bias towards A at both positions in all the samples analyzed (Figure 6).

The total number of different off-targets found across the 8 samples amounts to 34 (Table 2). Among those, 16 are uniquely found in one sample while 4 are found in 2 samples, 6 in 3 samples, 3 in 4 samples, 1 in 5 samples, 2 in 6 samples and 2 in 8 samples. The confidence in true editing events for off-targets is higher for the 18 sites that have been detected in more than one sample. However, we feel certain that even events detected in only one sample are true positive events for most of them because our threshold of detection is rather conservative (see Materials and Methods for details on our screening). Furthermore, their sequences generally fit the consensus logo defined across all samples. For example, the site 4471230, which is uniquely detected in sample 6, which co-expresses dsn3PLS-DYW with ORRM1 and RIP2 (Table 2), has a recognition sequence very similar to the consensus logo across all samples (Figure 6). The only noticeable discrepancy concerns the position -10 where a U is located in the sequence of the 4471230 site. There is no strong relationship between the level of editing extent of the off-targets and their detection in more than one sample (R^2^= 0.3). As an example, sites 433327 and 4471230 are edited at 22.9% and 21.2%, respectively, but are unique to sample 6 (Figure 6A, Table 2). Inversely, site 4350628 is detected in samples 2, 6, and 7 with a level of editing extent of 1.8%, 3%, and 2.9%, respectively (Table 2). This result comes from the contribution, in addition to the editing extent, of the JACUSA score, which depends on the quality of the reads and the depth of the coverage in the screening of the off-target events. The number of off-targets detected in each bacterial transcriptome assayed is strongly dependent on the level of editing extent of the *rpoA* target (Figure 7).

**Table 2.** Editing extent of the *rpoA* target and the off-target sites in the bacterial samples analyzed.

| sites<sup>a</sup> | sample1 | sample2 | sample3 | sample4 | sample5 | sample6 | sample7 | sample8 | sample9 | sample10 | sample11 | sample14 | #samples<sup>b</sup> |
| --- | --- | --- | --- | --- | --- | --- | --- | --- | --- | --- | --- | --- | --- |
| *rpoA* | **62.31<sup>c</sup>** | **71.0** | **30.2** | **57.3** | **59.1** | **81.9** | **78.5** | **68.0** | **58.1** | **22.7** | **15.6** | **4.5** | 12 |
| 301209 | **10.4** | **17.8** |  | 7.6 |  | **17.1** | **12.3** | 21.9 |  |  |  |  | 4 |
| 433327 | 5.4 |  |  |  |  | **22.9** |  |  |  |  |  |  | 1 |
| 779169 | **10.7** |  |  |  |  |  |  |  |  |  |  |  | 1 |
| 846395 | **1.2** | 0.7 |  | 0.2 |  | 1.1 |  | 0.9 |  |  |  |  | 1 |
| 957341 | 0.4 | **1.0** |  | 0.4 | 0.2 | 0.9 | 0.4 | 1.3 |  |  |  |  | 1 |
| 998460 | 5.2 |  |  |  | **10.9** | 3.2 |  |  |  |  |  |  | 1 |
| 1115962 | 0.8 | **1.1** |  | 0.7 | 0.5 | **2.7** | 1.1 | 1.6 | 0.6 |  |  |  | 2 |
| 1610233 | 3.8 |  |  |  |  | **25.0** |  | **17.5** |  |  |  |  | 2 |
| 1650242 |  |  |  |  |  | **2.3** | 0.6 |  |  |  |  |  | 1 |
| 1724437 | 6.5 | **13.7** |  |  | **14.8** | **18.2** |  |  |  |  |  |  | 3 |
| 1821466 | **38.5** | **14.9** |  | **10.9** |  | 25.0 | **28.0** | **46.7** | 20.0 |  |  |  | 5 |
| 1838803 | **14.6** | **10.1** | 0.9 | **6.8** | **6.0** | **20.3** | **11.1** | **18.3** | **9.4** |  | 0.4 |  | 8 |
| 1941416 | **16.8** | **21.8** |  | 6.0 | **14.3** | **37.4** | **26.6** | **33.9** | 9.3 |  |  |  | 6 |
| 1967867 |  | **10.8** |  |  |  | **24.0** | **14.7** |  |  |  |  |  | 3 |
| 2437684 | 4.4 | **14.0** |  |  |  | **18.6** | **11.3** |  |  |  |  |  | 3 |
| 2469966 | **5.3** |  |  |  |  | 7.5 |  |  |  |  |  |  | 1 |
| 2728082 | **6.1** | **2.2** |  | **5.0** |  | 1.3 | 0.8 | 0.9 |  |  |  |  | 3 |
| 2728396 | 0.8 | 1.5 | 0.8 | 0.8 | 1.6 | 2.0 | **2.0** | 2.0 | 1.0 | 1.0 | 0.6 | 0.1 | 1 |
| 2767599 | **7.5** | **6.0** |  | 5.7 | 2.4 | **32.9** | **12.5** | **13.4** | **10.1** |  |  |  | 6 |
| 3018155 | 1.8 | **13.2** |  |  |  |  |  |  |  |  |  |  | 1 |
| 3146210 | **2.1** | 1.4 |  | 1.0 | 0.5 | 1.3 | 0.3 | 1.4 | 0.6 |  |  |  | 1 |
| 3248710 | **11.1** | 3.5 |  | 5.3 | 4.2 | **46.6** | **18.2** | 9.1 | 5.7 |  |  |  | 3 |
| 3406026 | 1.8 | 1.5 |  | 1.0 | 0.9 | **6.9** | 2.3 | 2.6 | 1.0 |  |  |  | 1 |
| 3960072 | 3.7 | 3.4 |  | 1.4 | 1.8 | **14.1** | **8.7** | 4.1 | 2.7 |  |  |  | 2 |
| 4036064 | 0.6 | **1.4** |  | 0.5 | 0.7 | 1.1 | 0.5 | 1.2 | 0.6 | 0.3 |  |  | 1 |
| 4058658 | **1.4** | 0.6 |  | 0.8 | 0.4 | **2.1** | 0.6 | **2.0** | **1.1** |  |  |  | 4 |
| 4118470 | **9.7** |  |  |  |  |  |  |  |  |  |  |  | 1 |
| 4145999 | 1.0 | **1.5** |  | 0.8 | 0.4 | 0.7 | 0.4 | 1.2 | 0.4 |  |  |  | 1 |
| 4148094 | **7.3** | **5.1** |  | 4.7 | 3.7 | **14.1** | **8.2** | 8.1 | 3.2 | 0.9 | 1.0 |  | 4 |
| 4258541 | **38.1** | **41.6** |  | **16.5** | **46.8** | **41.5** | **45.2** | **45.7** | **23.2** |  |  |  | 8 |
| 4289975 | **10.0** |  |  |  |  | 21.9 |  |  |  |  |  |  | 1 |
| 4350628 | 0.6 | **1.8** |  | 0.4 | 0.2 | **3.0** | **2.9** | 0.3 |  |  |  |  | 3 |
| 4471230 |  |  |  |  |  | **21.2** |  |  |  |  |  |  | 1 |
| 4495686 | **5.8** | **5.4** |  | 2.6 | 4.9 | 7.1 | 7.4 | 4.4 |  |  |  |  | 2 |

<sup>a</sup>sites are denominated by their coordinated on the bacterial chromosome
<sup>b</sup>number of samples where the site was above the significant threshold for detection
<sup>c</sup>number in bold denote sites that passed the significant threshold

**Figure 7.**
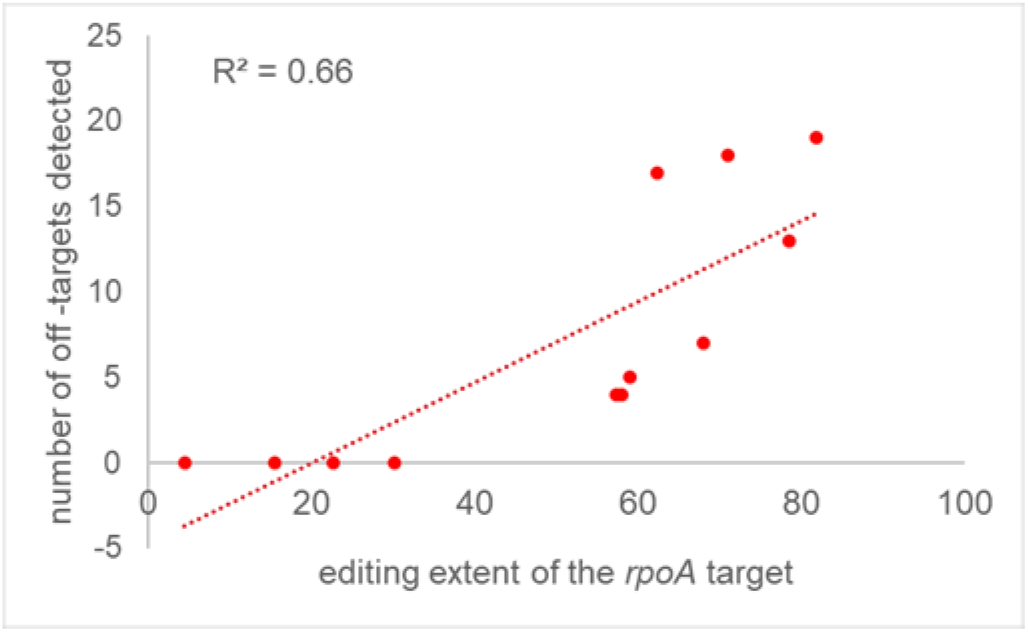
The number of off-targets detected in the bacterial transcriptome depends on the editing extent of the *rpoA* target. The level of editing extent of the *rpoA* target is a reflection of the efficiency of the editosome. The more efficient the editosome is, the higher number of off-targets.

Among the 34 off-target editing events, 21 occur in the annotated strand of coding sequence and result in an amino acid change (Supplementary Table S3). The highest edited off-target (21%), which results in a proline to leucine substitution, occurs in a hydroxyphenyl acetate permease. Interestingly, the two highest edited off-targets, 4258541 (37%) and 1821466 (26%), occur in the opposite strand to the annotated one. Because the bacterial genome is rather compact, these events might take place in the 5’ or 3’ UTRs of adjacent genes. We confirmed the existence of these two events by bulk-sequencing of RT-PCR products (Supplementary Figure S6). We chose two different samples to assay the level of editing extent of the two off-targets, sample1 (dsn3PLS-DYW + pCDF-RIP2) for 4258541 and sample 2 (dsn3PLS-DYW + pCDF-RIP9) for 1821466. Both samples show detectable level of editing with the 4258541 level of editing slightly lower (26% vs 38%) when compared to RNA-seq analysis, while 1821466 editing was slightly higher (20% vs 15%) when compared to RNA-seq analysis (Supplementary Figure S6, Supplementary Table S3).

## DISCUSSION

Except for RIP/MORF proteins, for which structures in complexes with a PPR editing factor have been obtained (19), there is little known about additional accessory proteins that affect editing extent in planta in corresponding mutants. The RIP/MORFs and ORRMs are common to chloroplasts and mitochondria, while OZ1 is specific to the chloroplast (12–14,20–23). We have taken three approaches to investigate the contribution of these accessory proteins in the editing process. Bacterial co-expression in the presence of the synthetic dsn3PLS-DYW PPR protein and the RNA target allowed us to test whether these proteins, either individually or in combination could influence the level of editing extent of the target. RNA electrophoretic mobility shift assays (REMSA) were used to measure the influence of the accessory proteins on the RNA binding activity of the synthetic factor. RNA-seq analysis of the bacterial transcriptomes of the different samples co-expressing the synthetic factor with various combinations of accessory proteins identified 34 off-targeting events. Investigation of their upstream sequences revealed both positions in adequation but also positions that diverge significantly from the PPR code.

Like Royan et al. (2021), we observed a positive contribution of RIP2 and RIP9 to the editing extent of the *rpoA*-C200 target by dsn3PLS-DYW in *E.coli*. However, the level of editing was significantly higher in our co-expression experiments. 61% for strains expressing RIP2 and 71% for RIP9 vs 35% (Figure 2). Several possibilities can explain this discrepancy: a different expression vector (pCDFDuet-1 vs pETM11 vs), a shorter version of RIP2 and RIP9 in the previous experiment, 48 aa (22 at N-terminus and 26 at the C-terminus) and 51 aa (15 at N-terminus and 36 at the C-terminus), respectively. These differences are significant, as RIP2 and RIP9 are relatively small proteins, so that the RIP/MORF proteins in Royan et al’s (2017) experiments lack ∼30% of the proteins’ compositions. In addition, in their experiments, one and two cysteines were mutated to serine, for RIP2 and RIP9 respectively, presumably to avoid the formation of disulfide bonds. In contrast to the prior report, we found that RIP9 contributes more to editing efficiency than RIP2 in two of the configurations we tested, with a tag either at the N or the C terminus.

We found that ORRM1, which had not previously been expressed in *E. coli* with a PPR editing factor and plant editing target, can result in around 30% editing of the RNA target. ORRM1 was originally identified by its homology to the RIP/MORF family (13). ORRM1 is an essential plastid editing factor; in Arabidopsis and maize mutants, RNA editing is impaired at particular sites, with an almost complete loss of editing for 12 sites in Arabidopsis and 9 sites in maize. The level of editing extent of *rpoA*-C200 in the Arabidopsis *orrm1* mutant is 26%. Therefore, ORRM1 is not essential to the editing of this site *in planta*, though its presence improves editing efficiency.

In contrast, OZ1 is essential to the editing of *rpoA*-C200 because editing is completely absent in the *oz1* mutant plant (14). Furthermore, although OZ1 has not been formally shown to interact with CLB19, these two proteins likely interact since OZ1 interacts with other components of the editing complex, such as OTP82 and CRR28 which are other PPR recognition factors (14). Therefore, we were surprised that we detected no effect of OZ1 in the bacterial expression system. One explanation may be that we predicted an incorrect transit sequence of 79aa, and perhaps the genuine sequence is shorter so that some important N-terminal sequence was absent in our expressed protein.

The lack of effect of ISE2 in the bacterial expression system may be due to its lower importance in *rpoA*-C200 editing. ISE2 is a chloroplast-localized RNA helicase that is required for multiple chloroplast RNA processing steps including RNA editing, splicing and processing of chloroplast ribosomal RNAs (15). The editing extent of *rpoA*-C200 is reduced to ca 40% in co-suppressed leaves (chlorotic tissues of mutant *ise2* plants expressing a 35S:ISE2–GFP transgene). Thus, ISE2 is not essential to the editing of *rpoA*-C200 *in planta*.

The co-expression of ORRM1 with RIP2 or RIP9 in the presence of the synthetic factor resulted in an editing extent of ∼80%, which is similar to the editing extent observed in the wild-type plants. The cumulative effect of these factors is almost perfectly additive suggesting that their involvement in the editing process is independent from each other. By contrast, RIP2 and RIP9 are redundant since their co-expression did not improve the editing extent of the target. This result is not really surprising since RIP2 and RIP9 are highly similar (60% identity, 74% similarity at the aa level). Furthermore, an AlphaFold prediction of dsn3PLS-DYW complexed with RIP2 or RIP9 shows a perfect alignment of these two latter proteins, suggesting that they bind the synthetic factor in a very similar way (Figure 8A, B, C).

**Figure 8.**
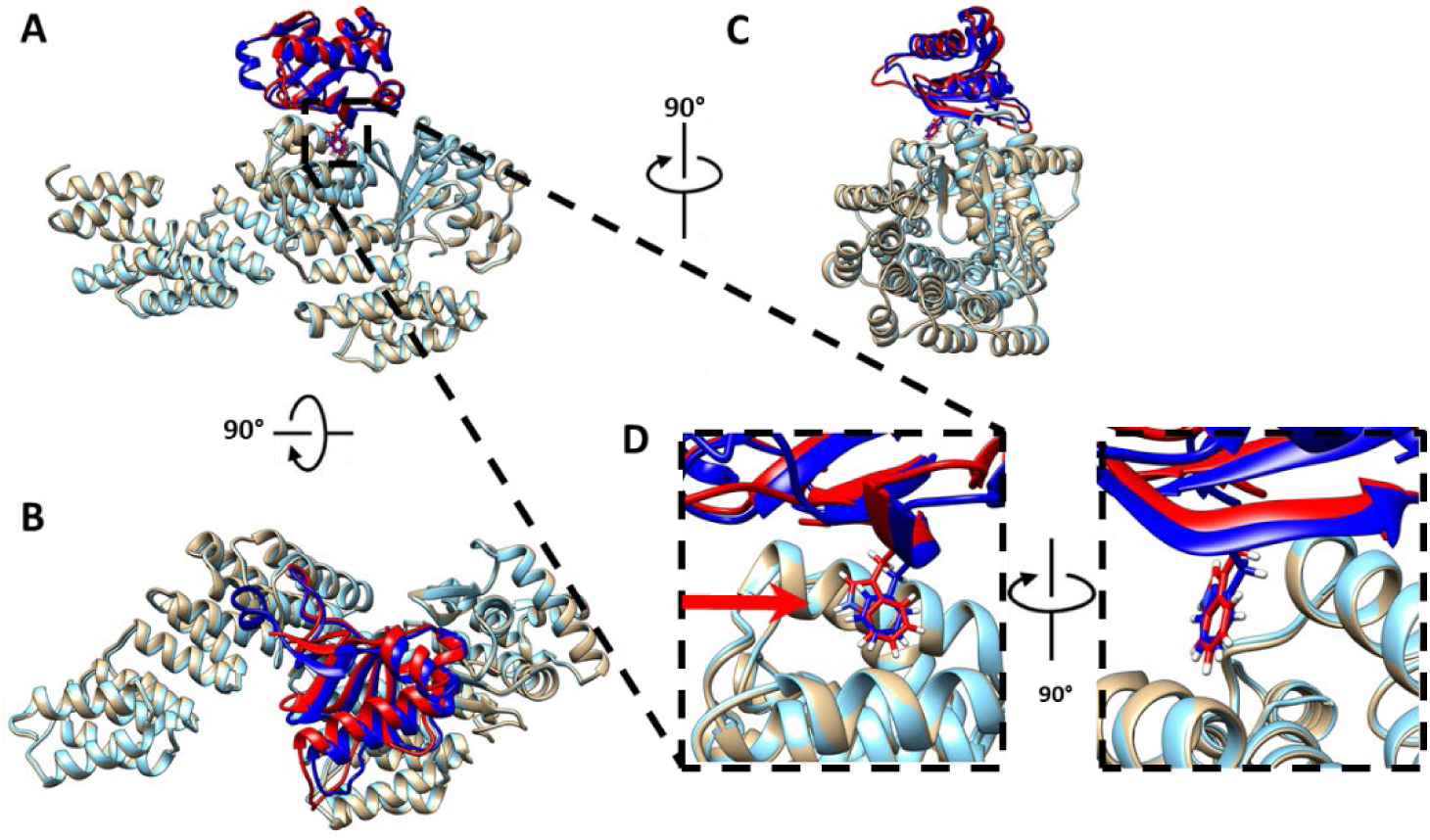
AlphaFold2-multimer predictions of RIP-dsn3PLS-DYW, superimposed, with each chain colour-coded. dsn3PLS-DYW (cyan), RIP2 (89–186) (blue), and RIP9 (86–192) (red). Three different views of the structures are shown, from side (**A**), top (**B**) and front (**C**). (**D**) Magnification depicting the RIP-PPR(L) predicted interaction. The arrow in panel D indicates the differential residue in RIP2 – Phe157 and in RIP9 – Trp160.

The gel shift assays indicate that the effects of RIP2 and RIP9 on the editing process are mediated by an interaction between these proteins and the PPR recognition factor that results in an enhanced binding activity of the factor for its RNA ligand. This increased RNA binding results from conformational change of the synthetic factor upon binding RIP9 (19). Our experiments also demonstrate that the editing efficiency in the bacterial expression system correlates with the RNA binding activity of the synthetic factor triggered by the accessory protein or the combination of them. For instance, RIP9 is significantly more efficient than RIP2 in editing of *rpoA*-C200 in bacteria in the presence of dsn3PLS-DYW (71% vs 61%). In REMSA, RIP9 also resulted in more protein complexed with RNA or smaller Kd, reflecting a better RNA binding activity of the synthetic factor for its RNA target (Figure 4). A closeup of the AlphaFold prediction of dsn3PLS-DYW complexed with RIP2 or RP9 may give us some clues to the reason of RIP9 better efficiency. At a position closest to the L repeat of dsn3PLS, a phenylalanine (157) is present in RIP2 while a tryptophan (160) is present in RIP9 (Figure 6D). This might explain the slightly better affinity of RIP9 and dsn3PL-DYW for *rpoA* in the REMSA.

Unexpectedly, we did not detect any gel shift when we assayed ORRM1 with target RNA (Figure 4B). In a previous study, ORRM1 fused to the maltose binding protein was shown to bind near several editing sites by REMSAs (13). However, we did not see any protein binding when our His-tagged ORRM1 was mixed with the RNA target or both the RNA target and the PPR editing factor (Figure 4). It is possible that purification of ORRM1 by nickel affinity has negatively affected ORRM1 activity. Alternatively, ORRM1 may have some other mode of action and does not influence the RNA binding activity of the PPR factor. Nevertheless, we showed that the presence of ORRM1 improves RIP2’s ability to enhance the RNA binding activity of the synthetic factor (Figure 5). This result reinforces the relationship between the editing efficiency observed in bacteria and the RNA binding affinity of dns3PLS-DYW. The higher the RNA binding affinity of the PPR factor for its target, the higher the editing extent in the bacterial expression system.

The pET-Duet expression system we used to co-express the synthetic factor and various combinations of accessory proteins in *E. coli* turned out to be a judicious choice in order to identify possible off-target events. This system, which is based on compatible vectors with different origin of replication and resistance makers, allowed the simultaneous and comparable expression of the synthetic editing factor and accessory proteins. As a result, we were able to identify a total of 34 different off-target editing events across all the bacterial samples analyzed.

The same synthetic factor and one of the accessory proteins assayed in this work, RIP2, had been assayed previously in bacteria (10). However, in this previous report no off-target events were detected. The authors proposed two explanations. First, the target transcript under the control of a strong T7 promoter is very abundant and may sequester a large fraction of the editing factor. This possibility also exists in our system. Second, the co-expression of RIP2 in their experiments reduced PPR protein expression by more than an order of magnitude. This drastic reduction of the expression of the synthetic factor when co-expressed with RIP2 was caused by the use of two vectors pETM20 and pETM11, respectively, that possess the same origin of replication. As a result, these two vectors competed for the expression machinery. Another factor possibly contributing to our increased detection of off-targets is that our depth coverage is 3 times higher than in the previous report (an average of 90 million mapped reads/sample, Supplementary Table S4).

The number of off-targets events identified in our study is comparable to what has been observed by expressing two moss editing factors in bacteria, PPR56 and PPR65 (9). RNA-seq transcriptome analyses after expression of the two editing factors revealed only seven off-targets for PPR65, most with editing efficiencies below 10%, but 79 sites of C-to-U editing for editing factor PPR56. However, the RIP-independent moss editing factors have reduced specificity because their L-motifs are unable to distinguish between different bases as the off-target sites generally show any of A, C, G or U aligned with the L-motifs in their proteins (9,24). The picture is quite different with dsn3PLS-DYW in the presence of accessory proteins, because the L motif at position-14 predominantly binds to the expected U based on the recognition code (Figure 6). This observation likely results from the interaction of the synthetic factor with either RIP2 or RIP9. Indeed, the first crystal structure of a designer PLS protein showed that L-motifs within this synthetic protein are slightly misaligned with the P and S motifs, but this misalignment is rescued by the formation of a complex with the cofactor RIP9 to allow for effective RNA recognition (19). The analysis of the off-target upstream sequences revealed other positions, such as -13, -9, -7, -6, which are in very good agreement with the PPR recognition code (Figure 6). The PPR code has been established by aligning PPR proteins with their target sequences and comparing the co-occurrence of aa in their fifth and last position with their associated nucleotides (25–27). The positions at -5 (T/R) and -4 (G/N) are neutral or slightly favored; nevertheless, there is a very strong bias towards A at these positions. In sample 6, which co-expresses the synthetic factor with ORRM1 and RIP2,18 of the 19 of the off-target upstream sequences contain an A at position -4 (Figure 6A). The same bias was observed in the off-targets identified in the chloroplast transcriptome after dsn3PLS-DYW was introduced in a *clb19* mutant background (10). Clearly, the PPR code needs to be refined to be able to explain this result.

Another limitation of the PPR code is the assumption of an absence of context in the favorability of the binding to a certain ribonucleotide. Clearly, this assumption is proved wrong in considering the L motif (P/D), which at position -14 binds preferentially U as predicted but at position-8 binds predominantly G (Figure 6A, B). Other positions deviating from the expected nucleotide based on the PPR code are -15 and -10. This type of data should be helpful for optimizing future designs by replacing the specificity determining aa at these positions by other combinations. Since the PPR code is degenerate, N/D or P/D are equally favorable to bind U (27). P/D could be tried at position-15 while N/D could be tried at position -8. However, modifying the specificity of a PPR motif can also have impact on the specificity of other PPR motifs specially at upstream positions, again emphasizing that the assumption of independence in the specificity of each PPR motif is oversimplistic (24). Other factors that can blur the predictability of editing events based on the PPR recognition code are the limited accessibility of targets by the editosome due to RNA secondary structure or protection by other RNA-binding proteins.

This work sheds light on the contribution of accessory proteins in the function of the angiosperm editosome. We have demonstrated the independent involvement of RIP2 (or RIP9) with ORRM1 in the activity of this molecular apparatus. Furthermore, the level of editing reached by co-expressing the synthetic factor with these two accessory proteins, around 80%, is comparable to what is naturally achieved in the chloroplast of *Arabidopsis thaliana*. Such a level of editing in a heterologous system has only been achieved by the moss editing factors. It remains to be seen whether testing this system in human cells, as has been done for the moss factor, will result in hundreds of off-targets as observed with the moss PPR proteins, or whether plant accessory proteins will affect the level of off-target editing.

## ACKNOWLEDGMENTS

The authors are very grateful for technical assistance by the RNA core facility at Cornell University.

## REFERENCES

1. Small, I.D., Schallenberg-Rudinger, M., Takenaka, M., Mireau, H. and Ostersetzer-Biran, O. (2020) Plant organellar RNA editing: what 30 years of research has revealed. Plant J, 101, 1040–1056.

2. Bentolila, S., Oh, J., Hanson, M.R. and Bukowski, R. (2013) Comprehensive high-resolution analysis of the role of an Arabidopsis gene family in RNA editing. PLoS Genet, 9, e1003584.

3. Mirdita, M., Schutze, K., Moriwaki, Y., Heo, L., Ovchinnikov, S. and Steinegger, M. (2022) ColabFold: making protein folding accessible to all. Nat Methods, 19, 679–682.

4. Okuda, K., Nakamura, T., Sugita, M., Shimizu, T. and Shikanai, T. (2006) A pentatricopeptide repeat protein is a site recognition factor in chloroplast RNA editing. The Journal of biological chemistry, 281, 37661.

5. Wagoner, J.A., Sun, T., Lin, L. and Hanson, M.R. (2015) Cytidine deaminase motifs within the DYW domain of two pentatricopeptide repeat-containing proteins are required for site-specific chloroplast RNA editing. J Biol Chem, 290, 2957–2968.

6. Hayes, M.L., Dang, K.N., Diaz, M.F. and Mulligan, R.M. (2015) A conserved glutamate residue in the C-terminal deaminase domain of pentatricopeptide repeat proteins is required for RNA editing activity. J Biol Chem, 290, 10136–10142.

7. Boussardon, C., Salone, V., Avon, A., Berthome, R., Hammani, K., Okuda, K., Shikanai, T., Small, I. and Lurin, C. (2012) Two interacting proteins are necessary for the editing of the NdhD-1 site in Arabidopsis plastids. Plant Cell, 24, 3684–3694.

8. Sun, T., Bentolila, S. and Hanson, M.R. (2016) The unexpected diversity of plant organelle RNA editosomes. Trends Plant Sci, 21, 962–973.

9. Oldenkott, B., Yang, Y., Lesch, E., Knoop, V. and Schallenberg-Rudinger, M. (2019) Plant-type pentatricopeptide repeat proteins with a DYW domain drive C-to-U RNA editing in Escherichia coli. Commun Biol, 2, 85.

10. Royan S, G.B., Colas des Francs-Small C, Honkanen S, Schmidberger J, Soet A, Sun Y, Vincis Pereira Sanglard L, Bond C, Small I. (2021) A synthetic RNA editing factor edits its target site in chloroplasts and bacteria.

11. Chateigner-Boutin, A.L., Ramos-Vega, M., Guevara-Garcia, A., Andres, C., de la Luz Gutierrez-Nava, M., Cantero, A., Delannoy, E., Jimenez, L.F., Lurin, C., Small, I., et al. (2008) CLB19, a pentatricopeptide repeat protein required for editing of rpoA and clpP chloroplast transcripts. The Plant Journal : for cell and molecular biology, 56, 590.

12. Takenaka, M., Zehrmann, A., Verbitskiy, D., Kugelmann, M., Hartel, B. and Brennicke, A. (2012) Multiple organellar RNA editing factor (MORF) family proteins are required for RNA editing in mitochondria and plastids of plants. Proc Natl Acad Sci U S A, 109, 5104–5109.

13. Sun, T., Germain, A., Giloteaux, L., Hammani, K., Barkan, A., Hanson, M.R. and Bentolila, S. (2013) An RNA recognition motif-containing protein is required for plastid RNA editing in Arabidopsis and maize. Proc Natl Acad Sci U S A, 110, E1169–1178.

14. Sun, T., Shi, X., Friso, G., Van Wijk, K., Bentolila, S. and Hanson, M.R. (2015) A zinc finger motif-containing protein is essential for chloroplast RNA editing. PLoS Genet, 11, e1005028.

15. Bobik, K., McCray, T.N., Ernest, B., Fernandez, J.C., Howell, K.A., Lane, T., Staton, M. and Burch-Smith, T.M. (2017) The chloroplast RNA helicase ISE2 is required for multiple chloroplast RNA processing steps in Arabidopsis thaliana. Plant J, 91, 114–131.

16. Xu, L., Liu, Y. and Han, R. (2019) BEAT: A Python Program to Quantify Base Editing from Sanger Sequencing. CRISPR J, 2, 223–229.

17. Pettersen, E.F., Goddard, T.D., Huang, C.C., Couch, G.S., Greenblatt, D.M., Meng, E.C. and Ferrin, T.E. (2004) UCSF Chimera--a visualization system for exploratory research and analysis. J Comput Chem, 25, 1605–1612.

18. Piechotta, M., Naarmann-de Vries, I.S., Wang, Q., Altmuller, J. and Dieterich, C. (2022) RNA modification mapping with JACUSA2. Genome Biol, 23, 115.

19. Yan, J., Zhang, Q., Guan, Z., Wang, Q., Li, L., Ruan, F., Lin, R., Zou, T. and Yin, P. (2017) MORF9 increases the RNA-binding activity of PLS-type pentatricopeptide repeat protein in plastid RNA editing. Nat Plants, 3, 17037.

20. Bentolila, S., Heller, W.P., Sun, T., Babina, A.M., Friso, G., van Wijk, K.J. and Hanson, M.R. (2012) RIP1, a member of an Arabidopsis protein family, interacts with the protein RARE1 and broadly affects RNA editing. Proc Natl Acad Sci U S A, 109, E1453–1461.

21. Hackett, J.B., Shi, X., Kobylarz, A.T., Lucas, M.K., Wessendorf, R.L., Hines, K.M., Bentolila, S., Hanson, M.R. and Lu, Y. (2017) An organelle RNA Recognition Motif protein is required for photosystem II subunit *psbF* transcript editing. Plant Physiol, 173, 2278–2293.

22. Shi, X., Castandet, B., Germain, A., Hanson, M.R. and Bentolila, S. (2017) ORRM5, an RNA recognition motif-containing protein, has a unique effect on mitochondrial RNA editing. J Exp Bot, 68, 2833–2847.

23. Shi, X., Hanson, M.R. and Bentolila, S. (2015) Two RNA recognition motif-containing proteins are plant mitochondrial editing factors. Nucleic Acids Res, 43, 3814–3825.

24. Lesch, E., Schilling, M.T., Brenner, S., Yang, Y., Gruss, O.J., Knoop, V. and Schallenberg-Rudinger, M. (2022) Plant mitochondrial RNA editing factors can perform targeted C-to-U editing of nuclear transcripts in human cells. Nucleic Acids Res, 50, 9966–9983.

25. Yagi, Y., Hayashi, S., Kobayashi, K., Hirayama, T. and Nakamura, T. (2013) Elucidation of the RNA recognition code for pentatricopeptide repeat proteins involved in organelle RNA editing in plants. PLoS One, 8, e57286.

26. Takenaka, M., Zehrmann, A., Brennicke, A. and Graichen, K. (2013) Improved computational target site prediction for pentatricopeptide repeat RNA editing factors. PLoS One, 8, e65343.

27. Kobayashi, T., Yagi, Y. and Nakamura, T. (2019) Comprehensive Prediction of Target RNA Editing Sites for PLS-Class PPR Proteins in Arabidopsis thaliana. Plant Cell Physiol, 60, 862–874.

