## supplemental material for "Accessory proteins increase the efficiency of RNA editing by Arabidopsis chloroplast editosomes"

### Slide 1
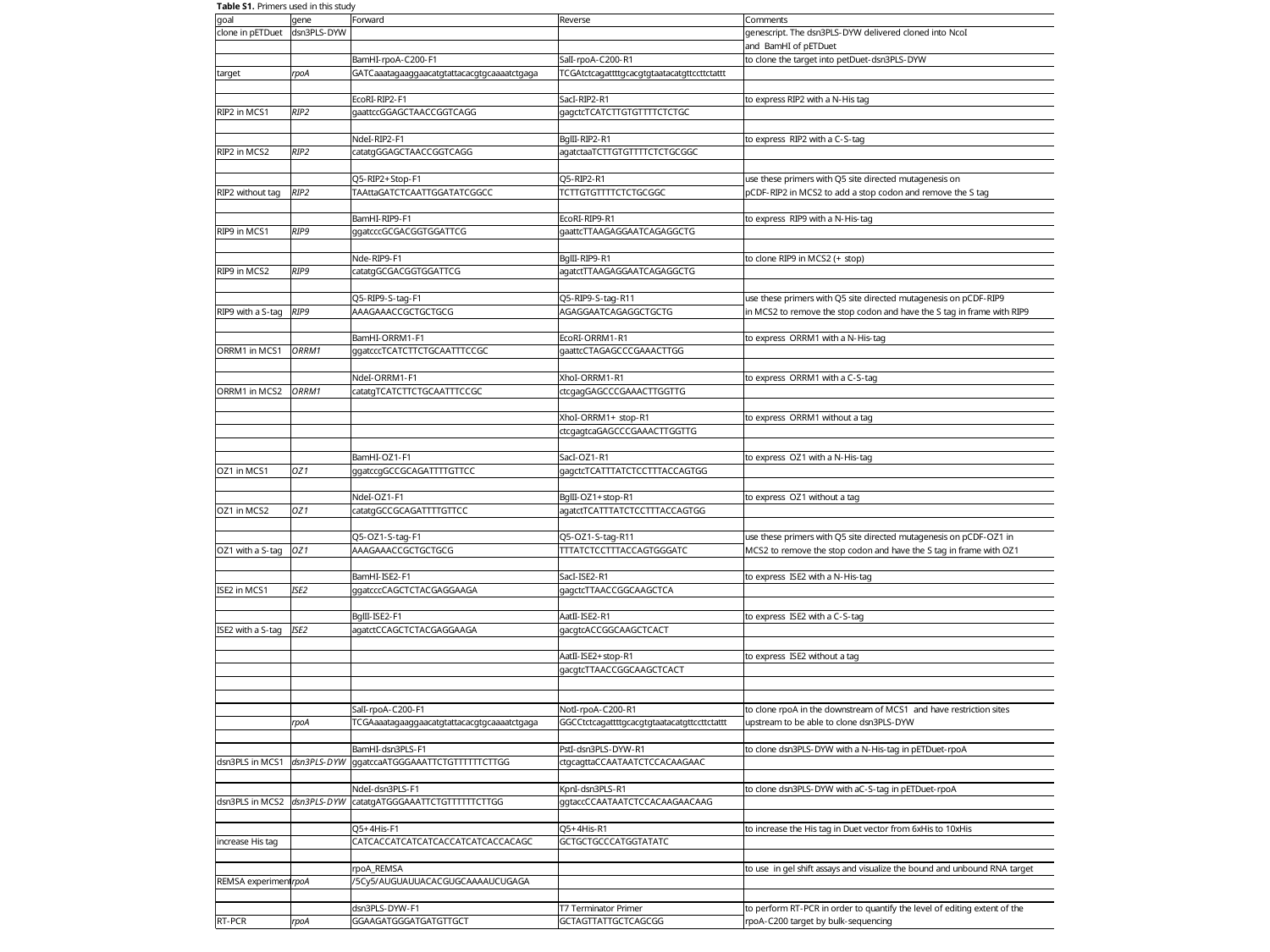

### Slide 2
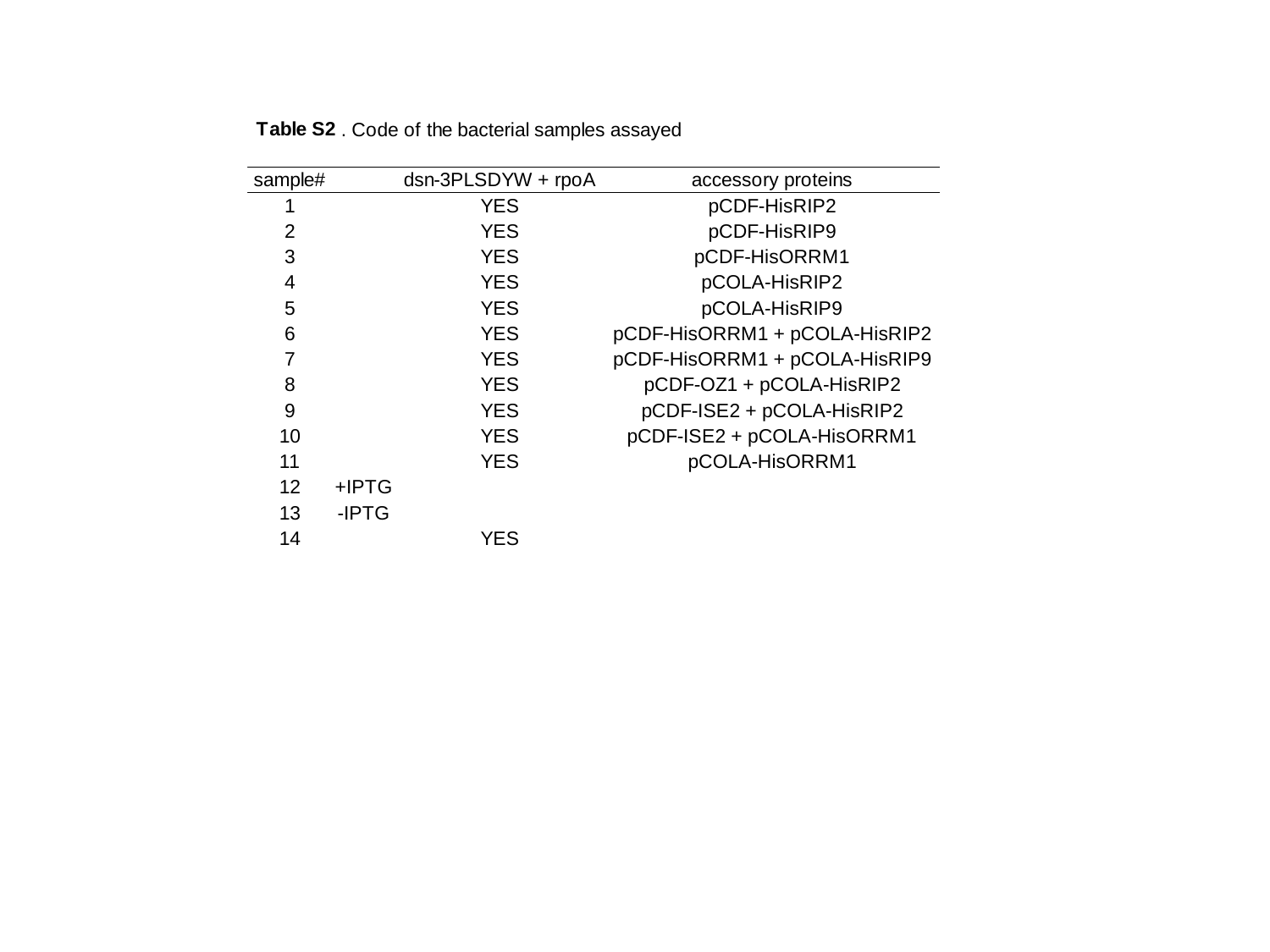

### Slide 3
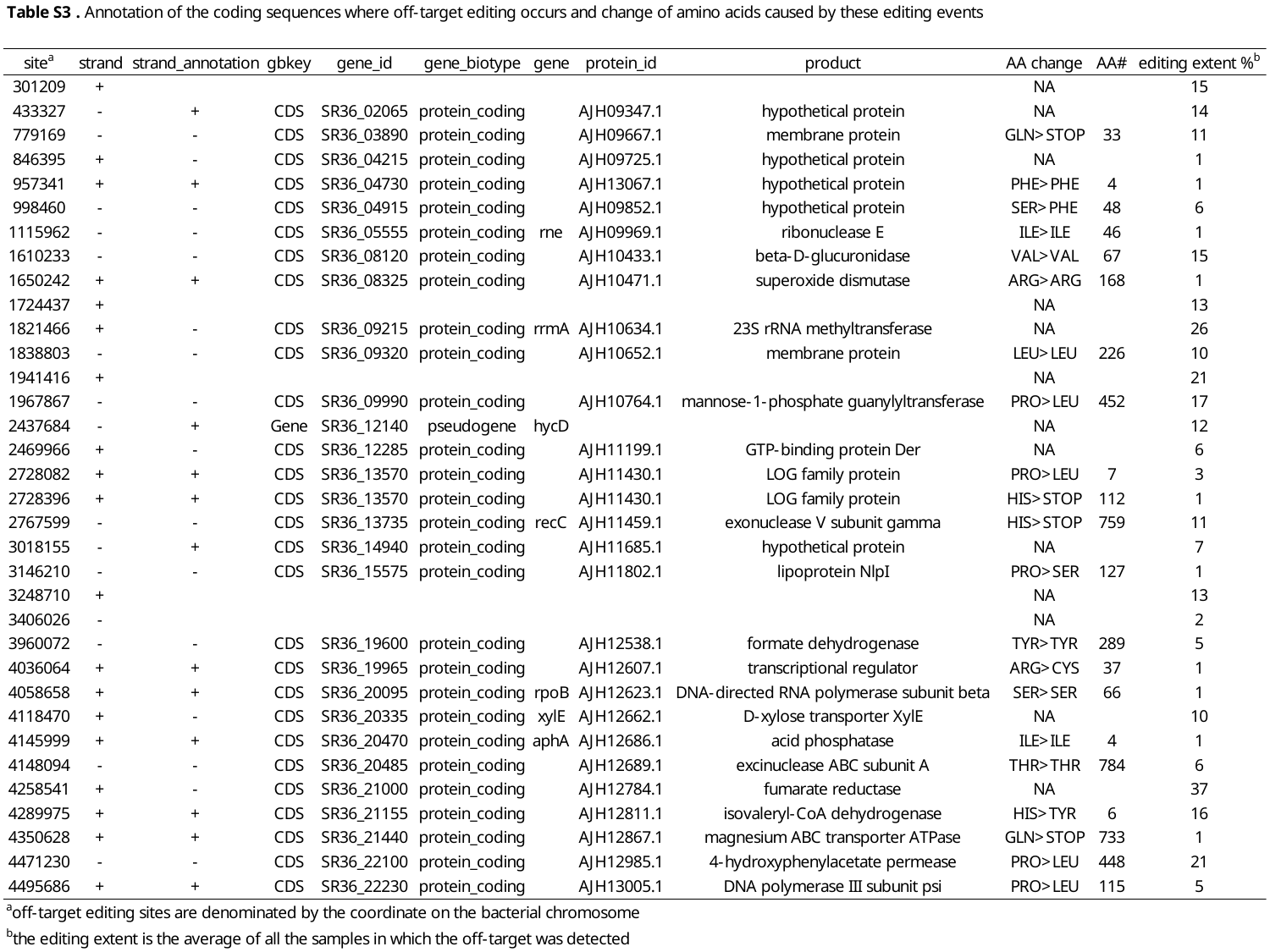

### Slide 4
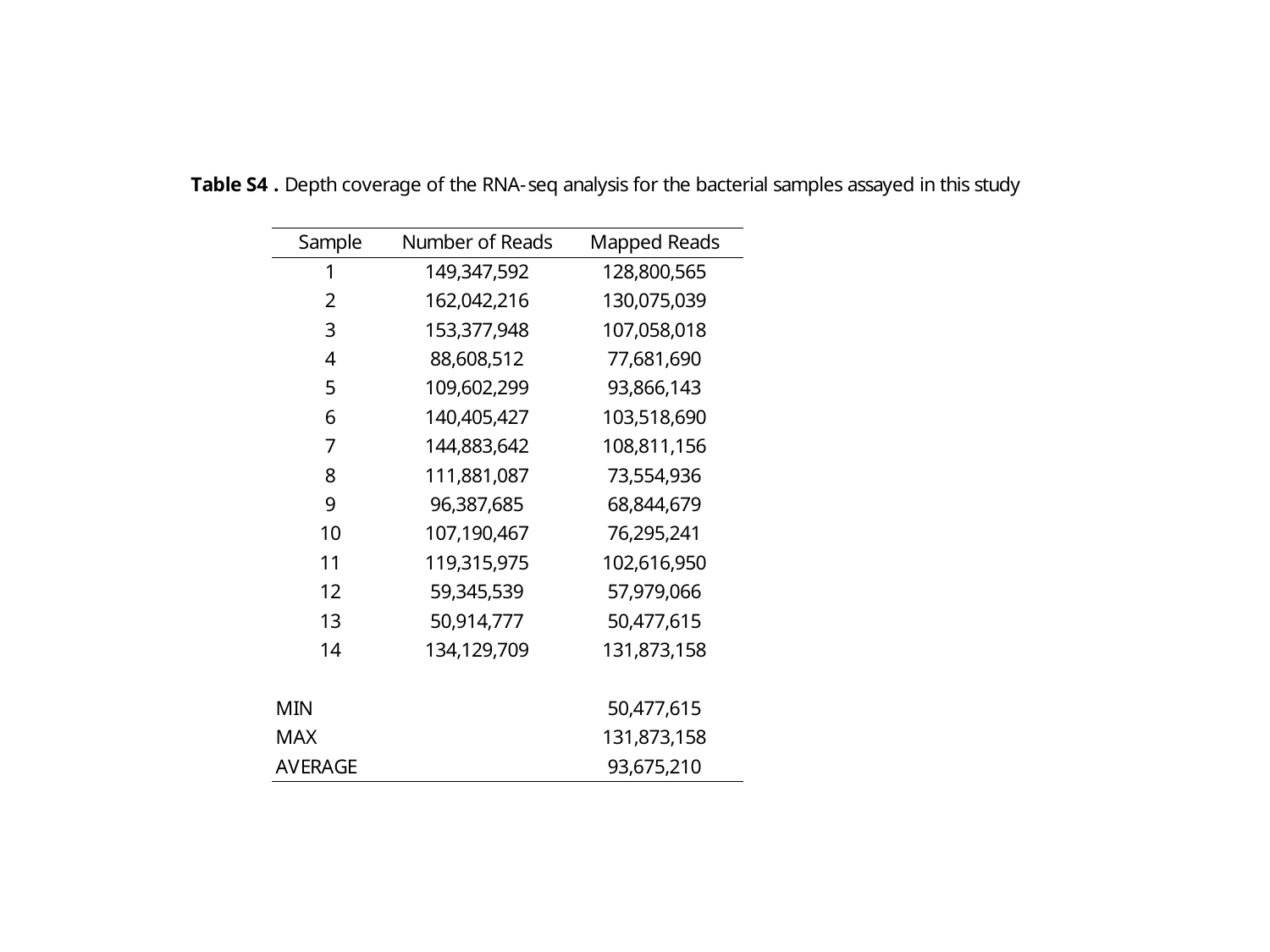

### Slide 5
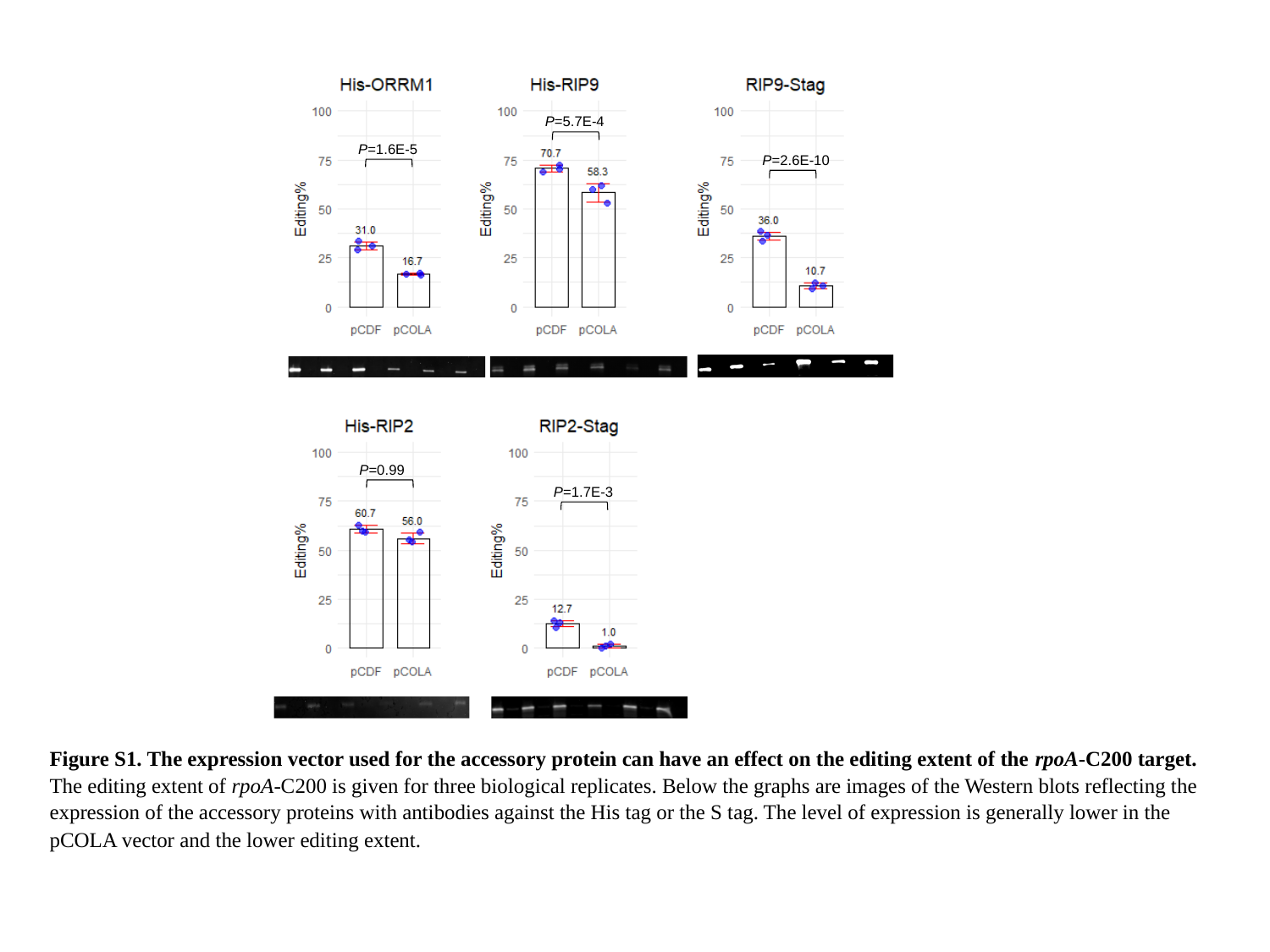

P=5.7E-4
P=1.6E-5
P=2.6E-10
P=0.99
P=1.7E-3
Figure S1. The expression vector used for the accessory protein can have an effect on the editing extent of the rpoA-C200 target. The editing extent of rpoA-C200 is given for three biological replicates. Below the graphs are images of the Western blots reflecting the expression of the accessory proteins with antibodies against the His tag or the S tag. The level of expression is generally lower in the pCOLA vector and the lower editing extent.

### Slide 6
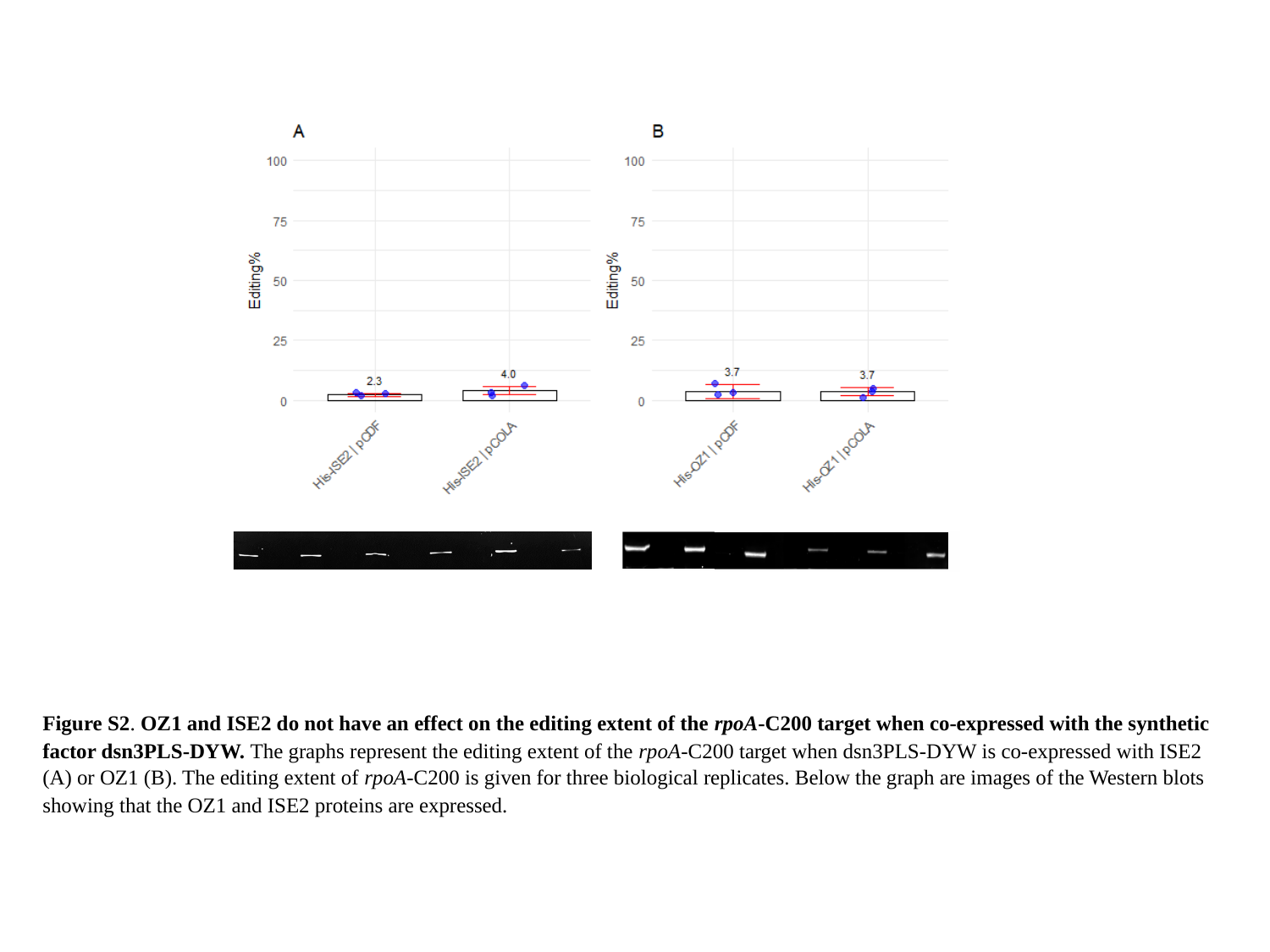

Figure S2. OZ1 and ISE2 do not have an effect on the editing extent of the rpoA-C200 target when co-expressed with the synthetic factor dsn3PLS-DYW. The graphs represent the editing extent of the rpoA-C200 target when dsn3PLS-DYW is co-expressed with ISE2 (A) or OZ1 (B). The editing extent of rpoA-C200 is given for three biological replicates. Below the graph are images of the Western blots showing that the OZ1 and ISE2 proteins are expressed.

### Slide 7
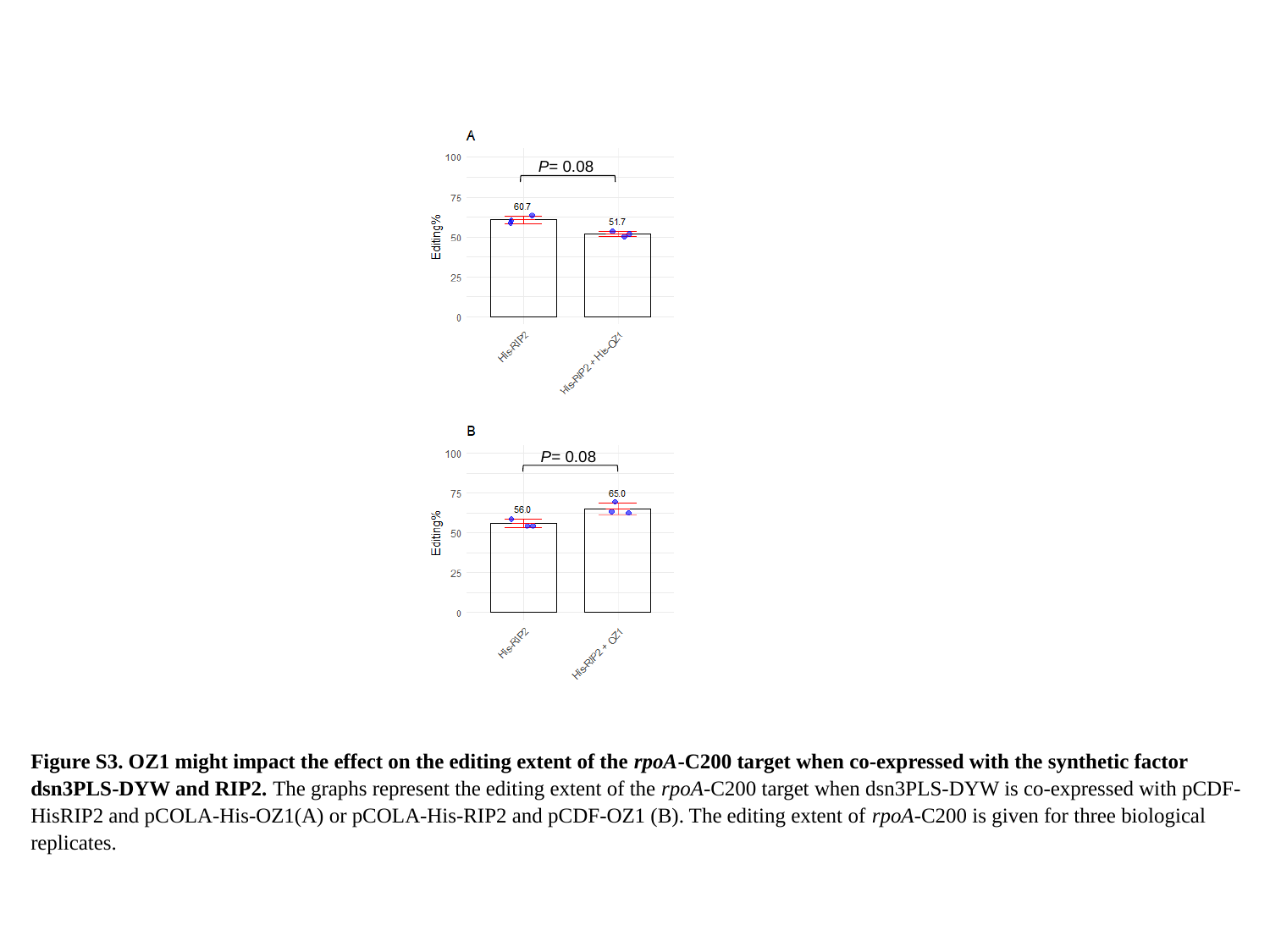

P= 0.08
P= 0.08
Figure S3. OZ1 might impact the effect on the editing extent of the rpoA-C200 target when co-expressed with the synthetic factor dsn3PLS-DYW and RIP2. The graphs represent the editing extent of the rpoA-C200 target when dsn3PLS-DYW is co-expressed with pCDF-HisRIP2 and pCOLA-His-OZ1(A) or pCOLA-His-RIP2 and pCDF-OZ1 (B). The editing extent of rpoA-C200 is given for three biological replicates.

### Slide 8
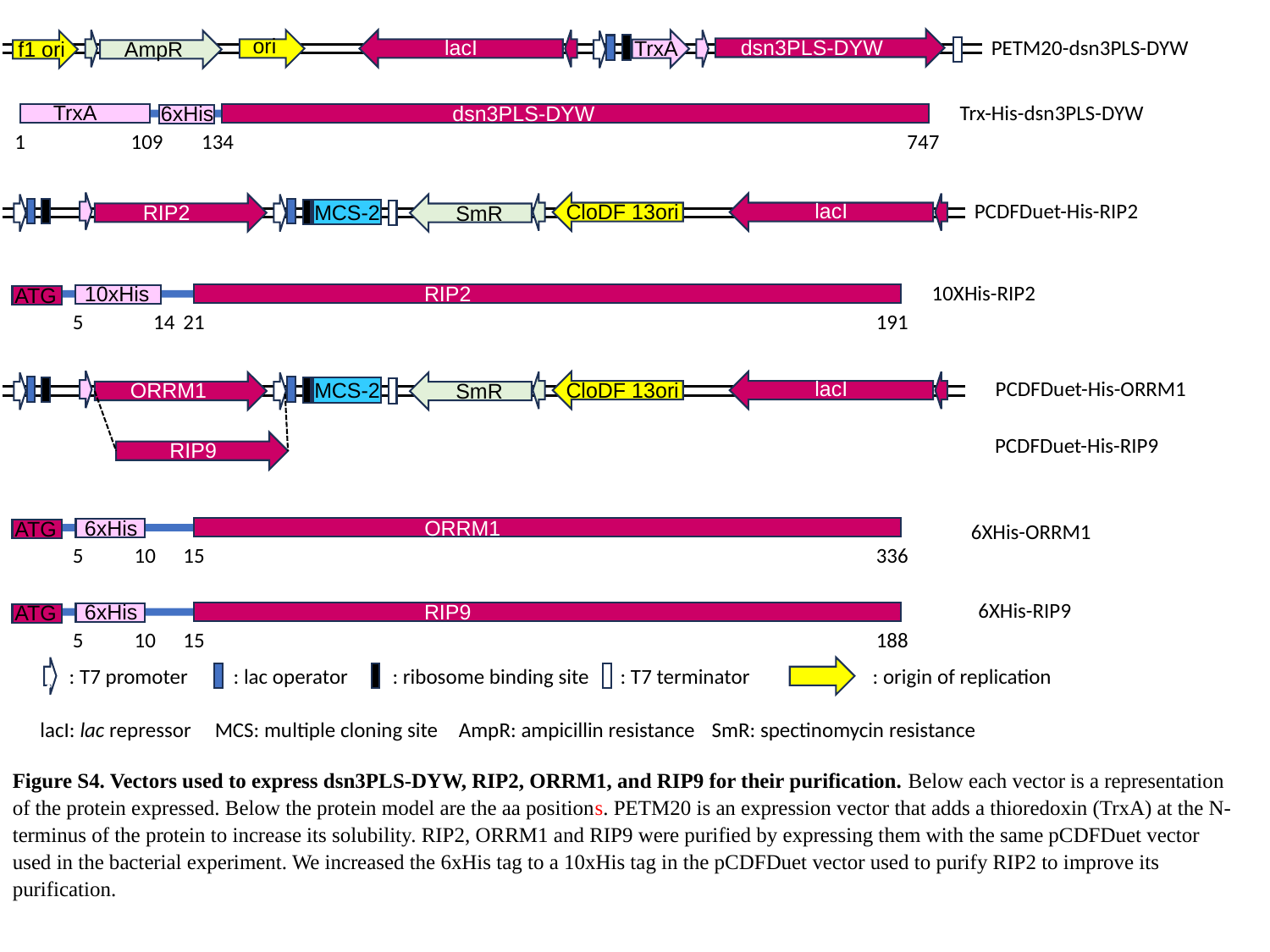

ori
PETM20-dsn3PLS-DYW
lacI
dsn3PLS-DYW
TrxA
f1 ori
AmpR
TrxA
Trx-His-dsn3PLS-DYW
6xHis
dsn3PLS-DYW
1
109
134
747
PCDFDuet-His-RIP2
lacI
CloDF 13ori
MCS-2
RIP2
SmR
10XHis-RIP2
10xHis
RIP2
ATG
5
14
21
191
PCDFDuet-His-ORRM1
lacI
CloDF 13ori
MCS-2
ORRM1
SmR
PCDFDuet-His-RIP9
RIP9
6xHis
ORRM1
ATG
5
10
15
336
6XHis-ORRM1
6XHis-RIP9
6xHis
RIP9
ATG
5
10
15
188
: T7 promoter
: lac operator
: ribosome binding site
: T7 terminator
: origin of replication
:
lacI: lac repressor
MCS: multiple cloning site
AmpR: ampicillin resistance
SmR: spectinomycin resistance
Figure S4. Vectors used to express dsn3PLS-DYW, RIP2, ORRM1, and RIP9 for their purification. Below each vector is a representation of the protein expressed. Below the protein model are the aa positions. PETM20 is an expression vector that adds a thioredoxin (TrxA) at the N-terminus of the protein to increase its solubility. RIP2, ORRM1 and RIP9 were purified by expressing them with the same pCDFDuet vector used in the bacterial experiment. We increased the 6xHis tag to a 10xHis tag in the pCDFDuet vector used to purify RIP2 to improve its purification.

### Slide 9
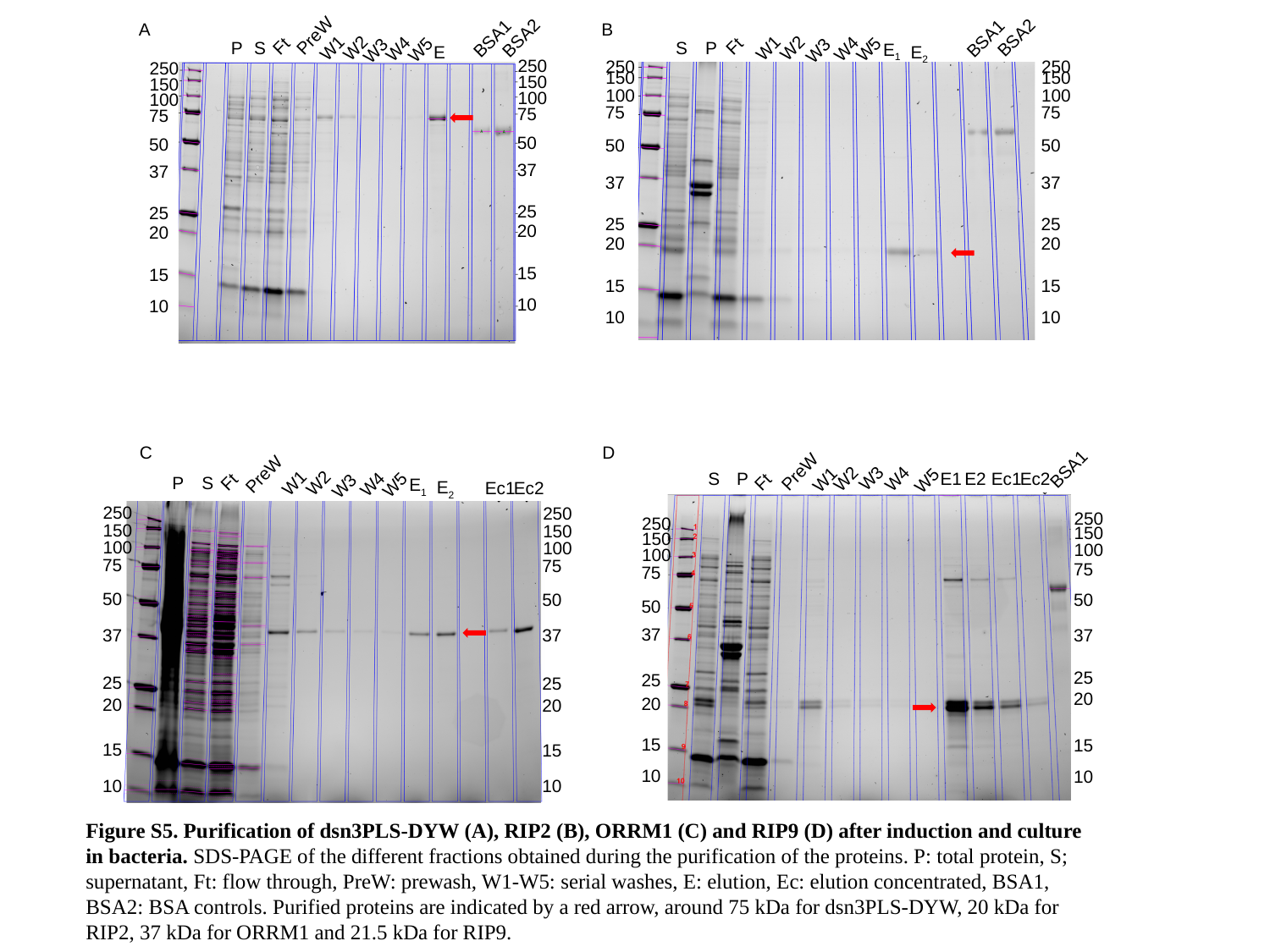

A
B
PreW
BSA2
BSA1
Ft
S
P
W1
W2
W4
W5
E_1_
W3
E_2_
BSA2
BSA1
Ft
P
S
W1
W2
W4
W5
W3
E
250
150
100
75
50
37
25
20
15
10
250
150
100
75
50
37
25
20
15
10
250
150
100
75
50
37
25
20
15
10
250
150
100
75
50
37
25
20
15
10
C
D
BSA1
PreW
PreW
W3
S
P
E1
E2
Ec1
Ec2
W2
W4
W1
W5
Ft
Ft
P
S
W1
W2
W4
W5
E_1_
W3
E_2_
Ec1
Ec2
250
150
100
75
50
37
25
20
15
10
250
150
100
75
50
37
25
20
15
10
250
150
100
75
50
37
25
20
15
10
250
150
100
75
50
37
25
20
15
10
Figure S5. Purification of dsn3PLS-DYW (A), RIP2 (B), ORRM1 (C) and RIP9 (D) after induction and culture in bacteria. SDS-PAGE of the different fractions obtained during the purification of the proteins. P: total protein, S; supernatant, Ft: flow through, PreW: prewash, W1-W5: serial washes, E: elution, Ec: elution concentrated, BSA1, BSA2: BSA controls. Purified proteins are indicated by a red arrow, around 75 kDa for dsn3PLS-DYW, 20 kDa for RIP2, 37 kDa for ORRM1 and 21.5 kDa for RIP9.

### Slide 10
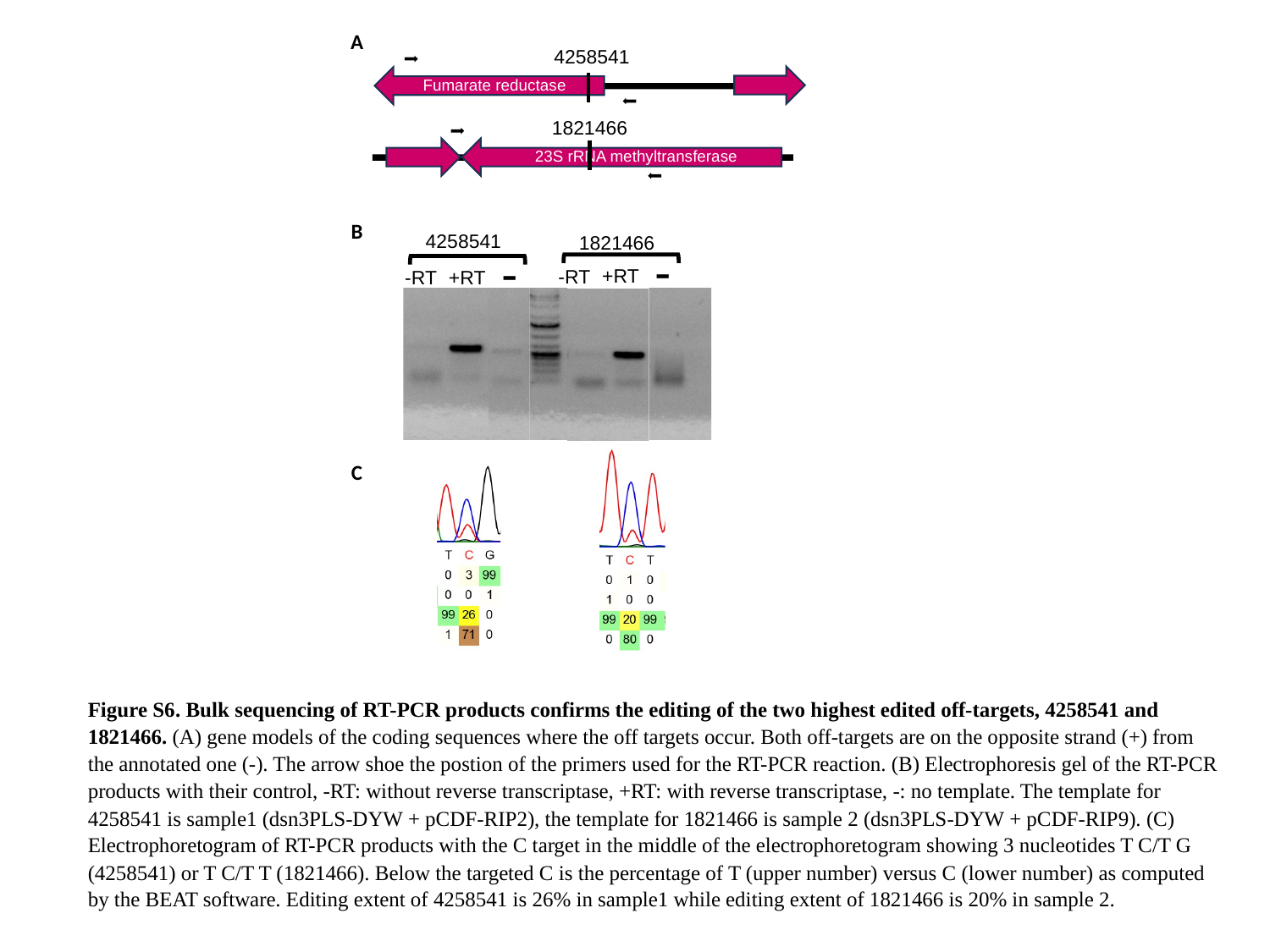

A
4258541
Fumarate reductase
1821466
23S rRNA methyltransferase
B
4258541
1821466
-
-
+RT
-RT
+RT
-RT
C
Figure S6. Bulk sequencing of RT-PCR products confirms the editing of the two highest edited off-targets, 4258541 and 1821466. (A) gene models of the coding sequences where the off targets occur. Both off-targets are on the opposite strand (+) from the annotated one (-). The arrow shoe the postion of the primers used for the RT-PCR reaction. (B) Electrophoresis gel of the RT-PCR products with their control, -RT: without reverse transcriptase, +RT: with reverse transcriptase, -: no template. The template for 4258541 is sample1 (dsn3PLS-DYW + pCDF-RIP2), the template for 1821466 is sample 2 (dsn3PLS-DYW + pCDF-RIP9). (C) Electrophoretogram of RT-PCR products with the C target in the middle of the electrophoretogram showing 3 nucleotides T C/T G (4258541) or T C/T T (1821466). Below the targeted C is the percentage of T (upper number) versus C (lower number) as computed by the BEAT software. Editing extent of 4258541 is 26% in sample1 while editing extent of 1821466 is 20% in sample 2.

### Slide 11
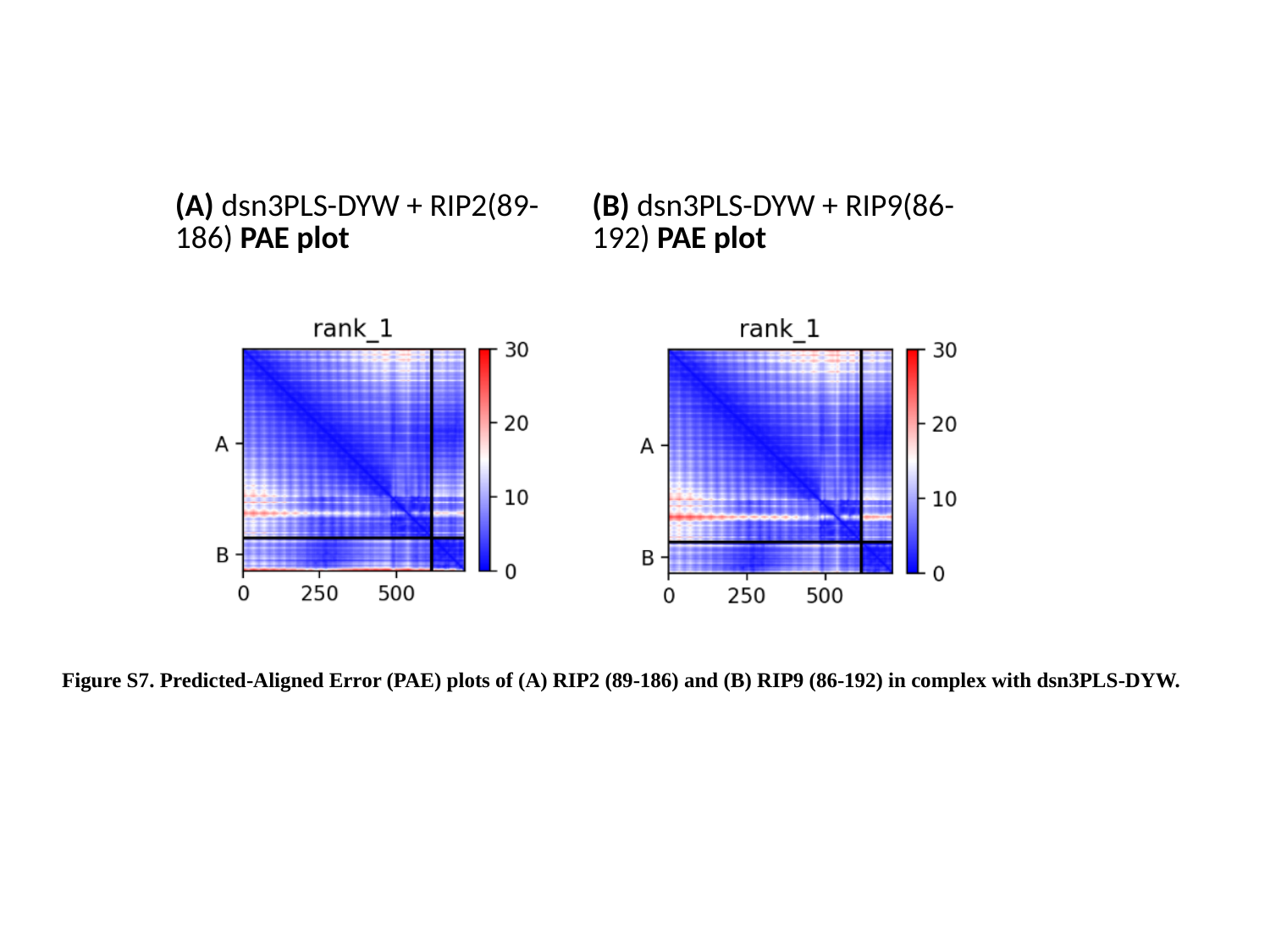

| (A) dsn3PLS-DYW + RIP2(89-186) PAE plot | (B) dsn3PLS-DYW + RIP9(86-192) PAE plot |
| --- | --- |
Figure S7. Predicted-Aligned Error (PAE) plots of (A) RIP2 (89-186) and (B) RIP9 (86-192) in complex with dsn3PLS-DYW.
